# Maternal iron deficiency perturbs embryonic cardiovascular development

**DOI:** 10.1101/2020.08.03.230615

**Authors:** Jacinta I. Kalisch-Smith, Nikita Ved, Dorota Szumska, Jacob Munro, Michael Troup, Shelley E. Harris, Aimée Jacquemot, Jack J. Miller, Eleanor M. Stuart, Magda Wolna, Emily Hardman, Fabrice Prin, Eva Lana-Elola, Rifdat Aoidi, Elizabeth M. C. Fisher, Victor L. J. Tybulewicz, Timothy J. Mohun, Samira Lakhal-Littleton, Eleni Giannoulatou, Duncan B. Sparrow

## Abstract

Congenital heart disease (CHD) is the most common type of birth defect, with a global prevalence of 0.9% of live births^1^. Most research in the last 30 years has focused on finding genetic causes of CHD. However, despite the association of over 100 genes with CHD, mutations in these genes only explain ~30% of cases^2^. Many of the remaining cases of CHD are caused by *in utero* exposure to environmental factors^3^. Here we have identified a completely new environmental teratogen causing CHD: maternal iron deficiency. In humans, iron deficiency anaemia is a major global health problem. 38% of pregnant women worldwide are anaemic^4^, and at least half of these are due to iron deficiency, the most prevalent micronutrient deficiency. We describe a mouse model of maternal iron deficiency anaemia that causes severe cardiovascular defects in her offspring. We show that these defects likely arise from increased retinoic acid signalling in iron deficient embryos, probably due to reduced activity of the iron-dependent retinoic acid catabolic CYP26 enzymes. The defects can be prevented by maternal iron administration early in pregnancy, and are also greatly reduced in offspring of mothers deficient in both iron and the retinoic acid precursor vitamin A. Finally, one puzzling feature of many genetic forms of CHD in humans is the considerable variation in penetrance and severity of defects. We show that maternal iron deficiency acts as a significant modifier of heart and craniofacial phenotype in a mouse model of Down syndrome. Given the high incidence of maternal iron deficiency, peri-conceptional iron monitoring and supplementation could be a viable strategy to reduce the prevalence and severity of CHD in human populations worldwide.

## Maternal iron deficiency perturbs embryonic development

Cardiac outflow tract (OFT) defects are the most common subtype of CHD^1^. In addition to arising from genetic mutation, such defects can be induced in mouse embryos by environmental factors including maternal exposure to hypoxia^5^ or by pharmacological activation of HIF1 signalling^6^. However, since such hypoxia studies do not replicate any particular clinical condition, it is uncertain how relevant these findings are to human populations. One way in which embryonic hypoxia might occur clinically is through maternal or embryonic anaemia. This is a major global health problem, affecting 20-40% of women of child-bearing age, a total of more than 500 million individuals^4^. Maternal anaemia in rabbits^7^ and maternal iron deficiency (ID) in rats^8^ can result in embryonic lethality, but the molecular mechanisms are unknown. In humans, at least half of all cases of anaemia result from ID^4^. Furthermore, low iron intake during pregnancy in humans may increase the risk of intrauterine growth restriction^9^ and CHD^10^. To investigate further, we used our previously published model of maternal ID^11^. Female C7BL/6J strain mice were weaned onto, and maintained continuously on, a low iron diet (2-6 ppm). At maturity, these mice had significantly reduced blood haemoglobin (Extended Figure 1a) and liver iron levels (Extended Figure 1b-e) compared to females fed a standard diet (200 ppm iron). ID females were mated to iron-replete C57BL/6J males, then maintained on the low iron diet until embryo collection on E15.5. ID embryo morphology was compared to that of embryos from control mothers (Figure 1a,b). Microscopic Magnetic Resonance Imaging (μMRI) showed that E15.5 embryos had significantly reduced liver iron levels (Extended Figure 1f-h). Macroscopic observation at dissection revealed that 12/80 ID embryos had died recently (0/58 controls, P=0.0010) and 26/68 of the surviving ID embryos had significant subcutaneous oedema (0/58 controls, *P*<0.0001). To determine the developmental progression of these phenotypes, we examined ID embryos between E9.5 and E14.5 (Figure 1c). Oedema and recent death were observed in a significant number of embryos from E12.5 onwards, with a peak at E13.5 (P=0.0008 vs E12.5 and P=0.0067 vs E15.5). Despite this, there was no significant difference in litter size (including dead embryos) between E10.5 and E15.5 (Extended Figure 1i), suggesting that additional undetected death and resorption of embryos was not occurring. We conclude that maternal ID severely perturbs embryonic development in mouse.

**Figure 1.**
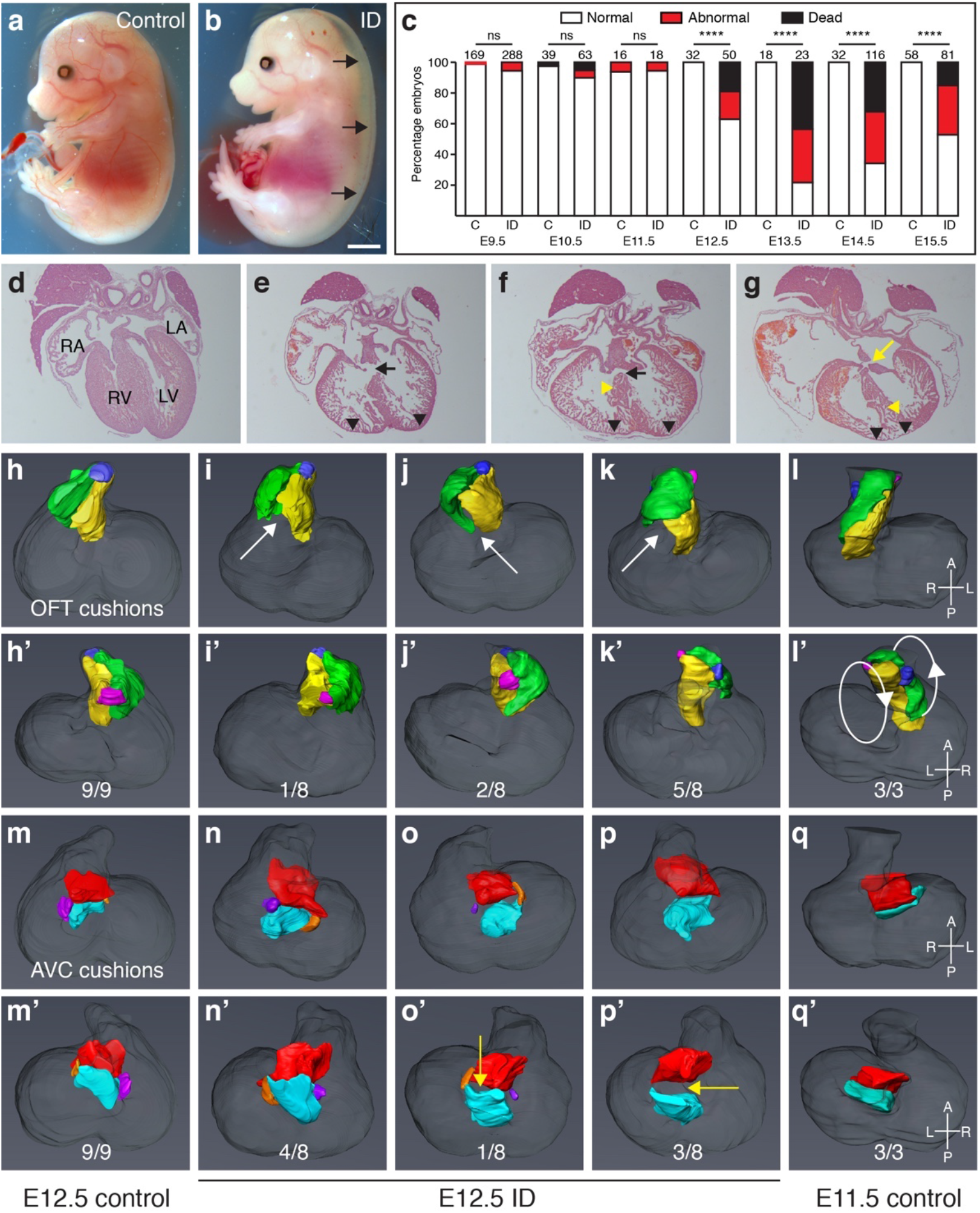
Maternal ID causes embryonic defects and lethality. (a-b) Embryos from ID mothers have gross sub-cutaneous oedema. Representative control (a) and ID (b) embryos. (c) Histograms of a developmental time-course showing significant embryonic lethality from E12.5. (d-g) Representative frontal H&E sections of hearts from control (d) and ID (e-g) E15.5 embryos. VSD (black arrows), AVSD (yellow arrow), thin ventricular myocardium (black arrowheads) and disorganised ventricular septum (yellow arrowheads) are indicated. (h-q) Abnormal cushion formation in E12.5 ID embryos. (h-l) Representative 3D Amira reconstructions of OFT cushions from manually-segmented HREM data showing dorsal (h-l) and ventral (h’-l’) views of (h) E12.5 control embryo, (i) normally-rotated OFT from E12.5 ID embryo, (j) partially-rotated OFT from E12.5 ID embryo, (k) non-rotated OFT from E12.5 ID embryo, (l) E11.5 control embryo. The left OFT cushion (yellow), right OFT cushion (green), aortic ICC (pink) and pulmonary ICC (blue) are shown. White arrows indicate unfused proximal cushions. (m-q) 3D reconstructions of the AV cushions showing dorsal (m-q) and ventral (m’-q’) views from the same embryos. The superior AV cushion (red), inferior AV cushion (cyan), left lateral AV cushion (orange) and right lateral AV cushion (purple) are shown. Yellow arrows indicate non-touching AV cushions. Scale bar = 2 mm (a,b) and 200 μm (d-g).

## ID embryos have cardiovascular defects

We hypothesised that maternal anaemia causes embryonic hypoxia, which we have previously shown leads to a variety of embryonic defects, most notably in the cardiovascular system^5^. To test this hypothesis, we examined cardiac morphology in surviving ID embryos at E15.5 (Figure 1d-g, Extended Table 1). 30/42 of these embryos had heart defects (1/37 controls, *P*<0.0001). Membranous and muscular ventricular septal defects (VSDs) were most common (Figure 1e,f), occasionally coupled with double-outlet right ventricle (DORV) or overriding aorta (OA). Finally, 5/42 embryos had atrioventricular septal defects (AVSD, Figure 1g, yellow arrow) compared to 0/37 controls (P=0.040). In addition, the ventricular myocardium was significantly thinner in all ID embryos (Figure 1e-g, black arrowheads; quantified in Extended Figure 2a), and in some embryos the ventricular septum (VS) was non-compacted (Figure 1f,g, yellow arrowheads). To investigate the developmental origins of the heart defects, we measured OFT physical parameters at E10.5 using 3D volume-rendered high resolution episcopic microscopy (HREM) datasets (Extended Figure 2b,c). Because OFT morphology changes rapidly, we matched embryos by Thieler stage (TS). In control embryos, distal OFT length increased from TS16 (30-34 somites) to TS 18 (40-44 somites), whilst the angle between distal and proximal OFT decreased. By contrast, at TS16 ID embryos had a significantly shorter distal OFT, and by TS18, they had a significantly larger distal/proximal OFT angle than controls. These observations are similar to mouse models of CHD caused by a reduced contribution of second heart field (SHF) cardiac progenitor cells to the OFT, including *Tbx1*, *Hes1* and *Hoxb1* null embryos, and embryos exposed to hypoxia *in utero*^5,12–14^. In these studies, reduced OFT length is proposed to cause malrotation and malalignment of the OFT at later developmental stages. In keeping with this hypothesis, by E12.5 5/8 ID embryos had a non-rotated OFT based on cardiac cushion position (Figure 1k), 2/8 were partially-rotated (Figure 1j) whilst only 1/8 was normal (Figure 1i), compared to 9/9 control embryos with normal OFT rotation (Figure 1h) P=0.0004). The non-rotated OFTs in ID embryos resembled those of control E11.5 embryos (Figure 1l).

We also found both OFT and atrio-ventricular (AV) cardiac cushions were abnormal at this stage. The cardiac cushions are the precursors of the valves and septa^15^. The OFT cushions form the aortic valve (AoV), pulmonary valve (PV) and the spiral septum dividing the OFT into the aorta and the pulmonary artery^16^. They are also required for the final closure of the tertiary ventricular foramen in the ventricular septum required to functionally separate the left and right ventricles^17^. The AV cushions contribute to the AV valves, as well the atrial and ventricular septa. In the OFT, the two major cushions are elongated and spiral around the OFT. In between the major cushions are two smaller ridges, called intercalating cushions (ICC) or intercalated valve swellings. By E12.5, the major cushions fuse to form the aortico-pulmonary septum as well as the right and left coronary cusps of the AoV and the right and left cusps of the PV. The ICC do not fuse: the aortic ICC forms the non-coronary (NC) leaflet of the AoV, and the pulmonary ICC forms the non-facing (NF) leaflet of the PV. At E12.5, proximal OFT cushion fusion had not occurred in 8/8 ID embryos (Figure 1i-k white arrows), compared to 2/9 controls (P=0.0019). Possibly this phenotype was due to generalised developmental delay, since ventricular volumes were significantly smaller in ID embryos (Extended Figure 2d). However, the appearance of external morphological landmarks was comparable between ID and control embryos, and by E13.5, the proximal OFT cushions were still not fused in 5/5 ID embryos (0/5 controls, P=0.0040). Concomitant with the morphological differences in the OFT at E12.5, the right OFT cushion and both ICCs were significantly smaller in ID embryos (Extended Figure 2f-h), although the left OFT cushion was not significantly changed (Extended Figure 2e). The aortic ICC was the most affected, being only 25% of normal volume, whilst the pulmonary ICC and right OFT cushion were ~70% normal size.

AV cushion formation was also abnormal in ID embryos. There are four AV cushions. The large inferior and superior cushions contribute to the atrial and ventricular septa, the aortic leaflet of the mitral valve (MV) and the septal leaflet of the tricuspid valve (TV). The smaller right lateral cushion forms the anterior and posterior leaflets of the TV, and the left lateral cushion forms the mural leaflet of the MV. At E12.5, the superior AV cushion volume was the same between control and ID embryos, but the inferior AV cushion was slightly smaller (Extended Figure 2i,j). In addition, the left and right lateral AV cushions were completely absent from 3/8 ID embryos (Figure 1p; 0/9 controls, P=0.0824), although there was no significant difference in volume between extant lateral cushions in ID embryos and controls (Extended Figure 2k,l). These cushions were also absent in 3/3 E11.5 control embryos (Figure 1q), suggesting that this might be due to delayed development. In addition, the inferior and superior cushions were not apposed in 4/8 ID embryos at E12.5, when fusion is normally complete (Figure 1o,p; yellow arrows; 0/9 controls, P=0.0294). These cushions were still not touching in 3/5 E13.5 ID embryos, suggesting a persistent defect rather than a developmental delay. The presence or absence of AV cushion contact did not correlate well with the extent of OFT rotation, as 3/5 embryos with the most severe OFT rotation defect (Figure 1k) had normal AV cushion contact, and 2/2 embryos with milder OFT rotation defects (Figure 1j) had non-touching AV cushions. In normal development, as the aortic root shifts from over the right ventricle to its final position over the left ventricle, the fused proximal OFT cushions align with the growing tip of the muscular VS^16,18^. The remaining interventricular communication is then closed at E14.5 by the membranous VS that forms by fusion of the rightward tips of the AV cushions with the proximal OFT cushions. Delayed or failed proximal cushion fusion, or incomplete shifting of the aortic root, will result in VSD, OA and DORV; and failed OFT-AV cushion fusion will prevent membranous VS formation and can contribute to AVSD. Thus, the cardiac defects observed at E15.5 in ID embryos are likely to arise from the OFT and AV cushion defects observed at E12.5.

Clinically, cardiac OFT defects commonly occur in conjunction with aortic arch (AA) anomalies. Typically, such defects originate from a failure of pharyngeal arch artery (PAA) formation or remodelling from E9.5. During normal development, the first three pairs of PAA form prior to TS16^19^. By TS16, the fourth pair appears, whilst the first two pairs regress, and no longer connect to the dorsal aorta (Figure 2a). The sixth pair develops by TS18, thus at this stage there are symmetrical arteries in the third, fourth, and sixth pharyngeal arches (Figure 2c). At TS16, whilst all ID embryos had fully patent PAA3, 5/12 ID embryos had bilaterally absent PAA4 (Figure 2b) and in the remainder, PAA4 was only partially formed on both sides, compared to 7/7 controls with fully patent PAA3-4, (P<0.0001). In addition, in 8/12 embryos the second PAA was still connected to the dorsal aorta (7 bilaterally and 1 unilaterally; Figure 2b’, black arrow), compared to 1/7 controls (Figure 2a,b; p=0.0399). By TS18, whilst all ID embryos had patent PAA3, 2/7 had absent or interrupted PAA4 (1 bilateral and 1 unilateral) and 3/7 had absent or interrupted PAA6 (2 bilateral and 1 unilateral; Figure 2d, white arrows; 6/6 controls with fully patent PAA3-6; P=0.1224). In addition, 4/7 ID embryos had bilaterally hyperplastic PAA2, with one of these still patent with the dorsal aorta (0/6 controls, P=0.049). This suggests that PAA development was delayed and abnormal. Indeed, at E12.5, 8/8 ID embryos had persistent right dorsal aorta (Figure 2f, orange arrow, 0/9 controls, P<0.0001). Unexpectedly, by E15.5 only 5/40 ID embryos had AA anomalies (Extended Table 2, 1/37 controls, P=0.1189). These included interrupted AA (IAA), aberrant right subclavian artery (A-RSA, Figure 2h, yellow arrow), right-sided AA and retroesophageal left subclavian artery (R-LCC, associated with right-sided AA). A similar decrease in the penetrance of AA anomalies between E10.5 and foetal stages has been observed in mouse knockout models^20,21^. Litter size did not differ significantly between E10.5 and E15.5 (Extended Figure 1i, P=0.7283), thus this did not result from increased lethality of embryos with abnormal AA formation prior to E15.5. This suggests that may be compensatory mechanisms allowing phenotypic recovery. The phenotypes observed at E15.5 are typical of a failure of PAA4 formation^22^. Therefore, the aortic arch defects in ID embryos are likely to arise from the observed perturbation of PAA4 formation earlier in development.

**Figure 2.**
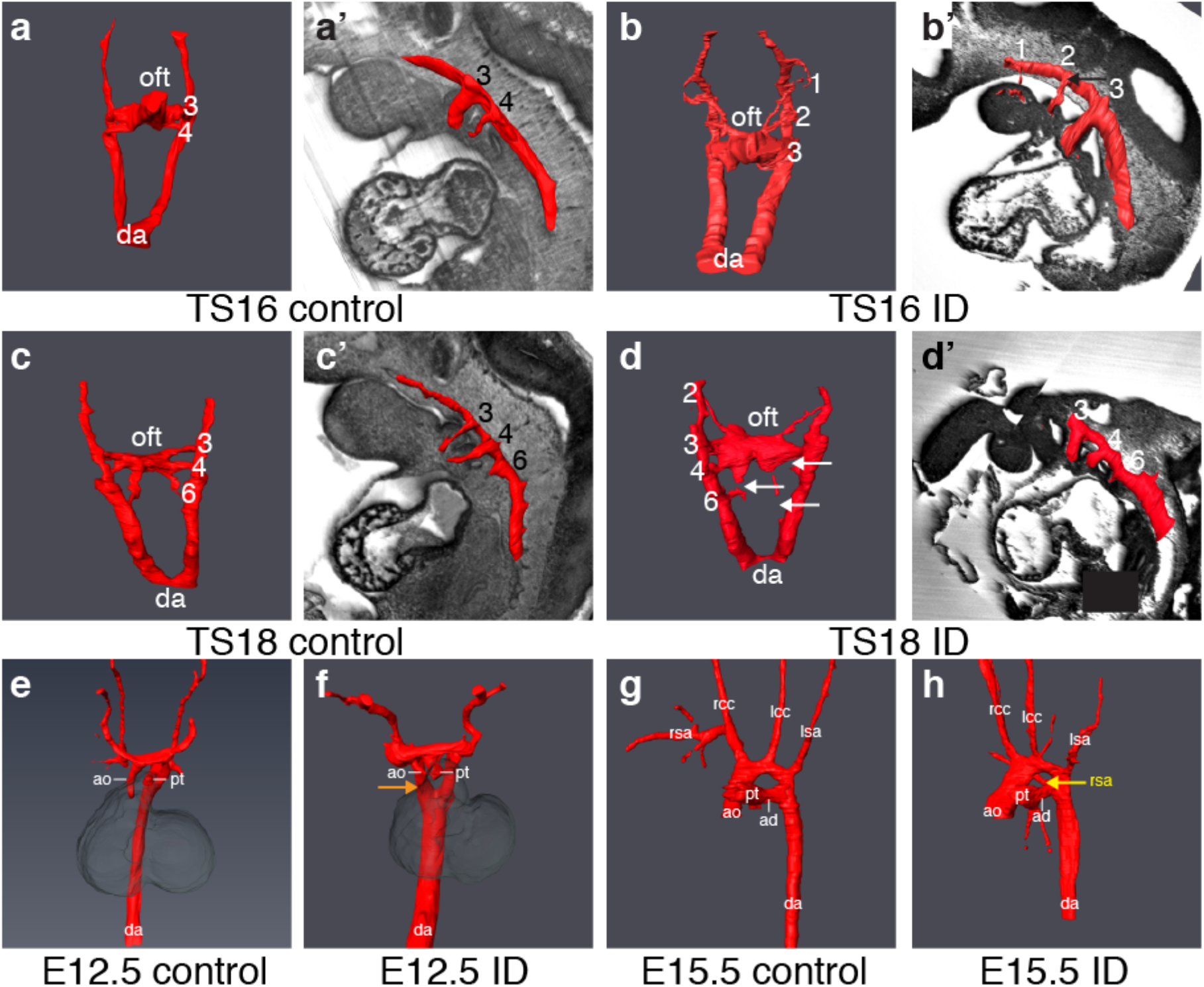
Maternal ID causes aortic arch abnormalities. Comparison of aortic arch artery morphology between control (a,c,e,g) and ID (b,d,f,h) embryos. Representative 3D Amira reconstructions from manually-segmented HREM data of early E10.5 (a,b), late E10.5 (c,d), E12.5 (e,f) and E15.5 (g,h) embryos. For clarity of PAA identity, 3D models of E10.5 embryos are overlaid onto a sagittal section of the same embryo (a’-d’). Persistent PAA2 (panel b’, black arrow), interrupted PAA4 and PAA6 (panel d, white arrows), persistent right dorsal aorta (panel f, orange arrow) and aberrant right subclavian artery (panel h, yellow arrow) are indicated. da = dorsal aorta; oft = outflow tract; ao = aorta; pt = pulmonary trunk; rsa = right subclavian artery; rcc = right common carotid artery; lcc = left common carotid artery; lsa = left subclavian artery; ad = arterial duct.

## Maternal ID causes premature differentiation of SHF cardiac progenitor cells

We next investigated the developmental origin of the cardiovascular defects. Four different cell lineages contribute to OFT cardiac cushion formation: endocardium, second heart field (SHF), cardiac neural crest (CNC) and epicardium^15^. Each particular cushion contains a characteristic combination of cells derived from a subset of these lineages. The aortic ICC was most severely effected in ID embryos (Extended Figure 2g). This cushion is predominantly composed of SHF-derived cells^23–25^, suggesting that the reduction in cushion size might be due to a lack of this cell type. A variety of mouse knockout models with cardiac OFT defects have a deficit of anterior SHF cells. This can arise by a variety of processes including premature differentiation^26^, disruption of proliferation^5,27,28^ or inhibited migration^29^. To investigate if any of these mechanisms were also occurring in ID embryos, we compared the transcriptomes of aSHF cells from ID and control embryos. We also compared these data with the transcriptome of aSHF cells from embryos exposed to hypoxia *in utero*, the molecular effects of which we have previously described^5^. ID and control female C57BL/6J mice were mated with hemizygous *Mef2c-AHF-GFP* males^30^. This allele directly drives GFP expression from the *Mef2c*-AHF enhancer element, specifically marking cells strongly in the aSHF and weakly in the distal OFT (Extended Figure 3a). For control and ID samples, embryos were collected at E9.5. For hypoxia samples, pregnant mice were exposed to an atmosphere containing 6% oxygen for four hours on E9.5, and embryos collected immediately after exposure. RNA was isolated from GFP+ cells from five individual somite-matched embryos for each condition and RNA sequencing (RNA-Seq) was performed. Unsupervised hierarchical clustering correctly grouped samples (Extended Figure 3b). As expected from our previous study^5^, differential expression (DE) analysis showed that aSHF cells from hypoxic embryos had increased expression of unfolded protein response (UPR) genes and spliced *Xbp1* transcript; elevated transcript levels of hypoxia-response genes (including those involved in metabolism, angiogenesis and pathway regulation); increased expression of cell cycle inhibitors; decreased expression of cell-cycle progression genes; but no induction of apoptotic HIF1 targets (Extended Figure 3d). By contrast, aSHF cells from ID embryos did not activate UPR response genes or HIF1 targets, nor was cell-cycle gene expression altered. Overall, there was very little overlap in the significantly DE genes between ID and hypoxia samples (Extended Figure 3c). We used gene ontology (GO) analysis with the GSEA online resource^31,32^ to provide an unbiased assessment of significantly altered pathways. As expected, the top upregulated gene set in the hypoxia samples was Hallmark_hypoxia (P=5.33×10^−55^), and the top downregulated gene set was Hallmark_mitotic_spindle (P=7.66×10^−10^). By contrast, the most enriched gene set in the ID model was Hallmark_myogenesis (P=4.65×10^−17^). Further analysis of the ID dataset revealed that expression of multiple markers of cardiac OFT differentiation were significantly upregulated (Extended Figure 3d). We also compared the top significantly DE transcripts from ID embryos with data from single cell transcriptomic analysis of the cardiogenic regions of E9.25 embryos^33^. The ID transcriptome was most closely related to the OFT cluster, rather than to the aSHF cluster (P=3×10^−6^, Extended Figure 3e).

Premature differentiation of the aSHF is a feature of several mouse knockout models with similar cardiac defects to our ID model, including *Tbx1* null^26,34^ and *Hoxb1* null^14^ embryos. In these models, premature differentiation of aSHF cells at E9.5 causes similar phenotypes to ID, namely defective OFT elongation, alignment and septation, resulting in a specific set of cardiac defects by E15.5 including membranous VSDs (with or without DORV or OA), transposition of the great arteries (TGA) and persistent *truncus arteriosus* (PTA), as well as AA anomalies. We validated the presence of premature differentiation in the SHF by examining protein expression levels and pattern of the myocyte differentiation marker MHC (Figure 3a-e). MHC expression is normally restricted to the wall of the OFT, with no expression in the contiguous cells of the aSHF (Figure 3a). By contrast, in ID embryos ectopic MHC expression extended into the aSHF (Figure 3b), and a significantly larger number of aSHF cells expressed MHC. Similar defects can be caused by reduced cell proliferation^35,36^ or activation of cell death^35^ in the SHF. However, there was no significant difference in the percentage of phosphorylated histone H3-positive nuclei between ID and control SHF cells (Extended Figure 4a-d), and no apoptosis was detected in the SHF by active/pro-CASPASE 3 staining (data not shown). The cardiac transcription factor GATA4 is a key regulator that promotes the switch from proliferation to differentiation in SHF cells^26^. *Gata4* transcripts were significantly increased in the ID aSHF transcriptome (Extended Figure 3d), and GATA4 protein levels were ectopically expressed in the aSHF of ID embryos at E9.5 (Figure 3f-i). By contrast, the expression levels and domain of TBX5 protein, a marker of the posterior SHF, were unchanged (Extended Figure 4e-g). GATA4 directly activates the transcription of many cardiac differentiation genes and transcription factors, including MHC^37^. Many of these direct GATA4 targets were also significantly upregulated in the ID aSHF transcriptome (Extended Figure 3d). In *Tbx1* null embryos, *Gata4* expression is elevated, and SHF cells prematurely differentiate^26^. Thus, it is likely that the cardiovascular defects in the ID embryos result from premature differentiation of aSHF cells, which then fail to migrate into the OFT, OFT cushions and aortic arches.

**Figure 3.**
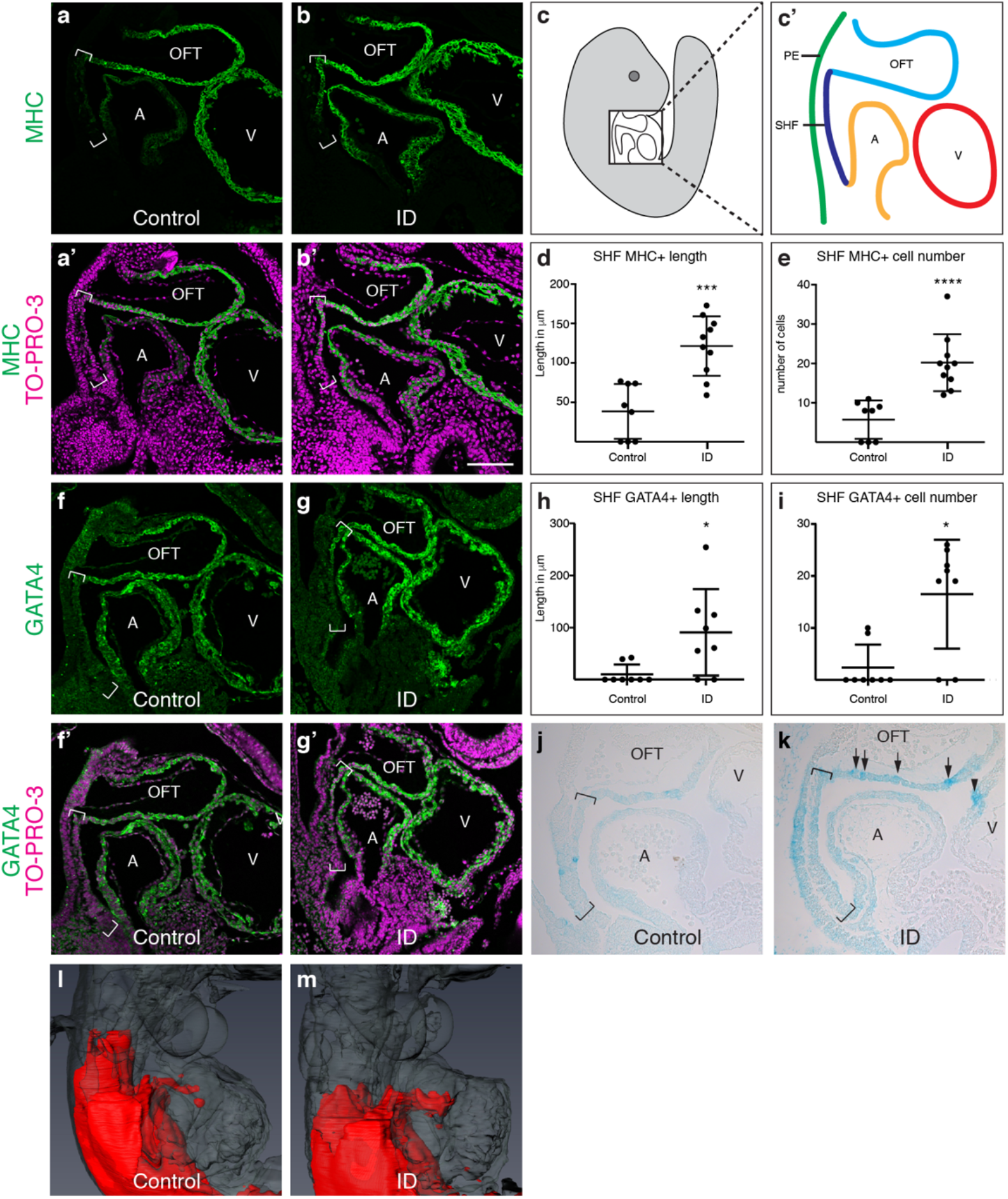
Effects of ID on cardiac progenitors and RA signalling. (a-e) Comparison of expression levels of the myocyte differentiation marker MHC (green) in sagittal sections of control (a,a’) and ID (b,b’) E9.5 mouse embryos by immunohistochemistry. Nuclei were stained with TO-PRO-3 (magenta). Location of the SHF is indicated by brackets. (c,c’) Diagrams indicating the relative positions of the SHF (dark blue), pharyngeal endoderm (green), OFT (light blue) and left ventricle (V, red) and left atrium (A, orange) in a sagittal section of an E9.5 embryo. Quantification of length of MF20-positive SHF (d) and number of MHC-positive cells in the SHF (e). (f-h) Comparison of expression levels of GATA4 (green) in control (f,f’) and ID (g,g’) E9.5 mouse embryos by immunohistochemistry. Nuclei were stained with TO-PRO-3 (magenta). Location of the SHF is indicated by brackets. (h) Quantification of length of GATA4-positive SHF. (j-m) Comparison of RA signalling levels in control (j,l) and ID (k,m) E9.5 embryos carrying the RARE-LacZ reporter allele. (j,k) Sagittal sections of E9.5 embryos stained with X-gal in wholemount. Location of the SHF is indicated by brackets. Increased and patchy staining in the OFT (arrows) and increased staining in the ventricle (arrowhead) are indicated. (l,m) Representative 3D Amira reconstructions of X-gal staining (red) from automatically thresholded two-channel HREM data derived from E9.5 embryos stained in wholemount. The embryos are shown in transparent grey. *** P<0.001, **** P<0.0001. Scale bar = 130 μm (a,b,f,g) and 95 μm (j,k).

## Embryos show perturbed retinoic acid signalling

The cardiovascular defects observed in ID embryos resemble those present in mouse or chick embryos with excess retinoic acid (RA) signalling^38–40^, after exposure to excess vitamin A^41^, or in humans exposed to the drug isotretinoin^42^. In mouse embryos, RA signalling is mediated by all-*trans* retinoic acid (ATRA). This compound is synthesised from vitamin A, and can be subsequently catabolised to the biologically inactive compound 4-hydroxy ATRA by the CYP26 family of cytochrome P450 enzymes. The active site of CYP26 enzymes incorporates haem, so we hypothesised that RA catabolism might be reduced in ID embryos, resulting in excess RA signalling. Supporting this hypothesis, *Cyp26b1* null mouse embryos have a similar spectrum of heart defects to ID embryos (C. Roberts, personal communication). In addition, *Gata4* transcription is directly activated by RA signalling^43,44^, thus this might also explain the ectopic upregulation of *Gata4* in ID embryos. To test this hypothesis, we examined the level of RA signalling in ID embryos using the RARE-LacZ transgenic reporter^45^. Males carrying the RARE-LacZ allele were crossed with control or ID C57BL/6J females, embryos collected at E9.5, and stained for ß-galactosidase activity with X-gal. In control embryos at E9.5, ß-galactosidase activity was present in paraxial mesoderm, SHF, OFT and brain, as previously described^45^. ID embryos had a broadly similar expression pattern, however X-gal staining developed more rapidly, suggesting higher levels of RA signalling in ID embryos. Sections from embryos dissected on the same day and stained for the same time confirmed that ID embryos had stronger SHF and OFT expression than controls (Figure 3j,k). In addition, expression was patchy in ID embryos (Figure 3k, arrows) and was also present in the ventricle (Figure 3k, arrowhead). Two-channel HREM of X-gal-stained E9.5 embryos coupled with 3D reconstruction in Amira was used better visualise the 3D pattern of RA expression in the OFT (Figure 3l,m). This shows that RA signalling levels were present in a larger domain of the OFT. These data suggest that ID embryos have increased RA signalling in the SHF and OFT, causing ectopic activation of *Gata4* in the SHF and initiating premature differentiation of these cells.

## Other embryonic processes dependent on RA signalling are perturbed in ID embryos

In addition to cardiovascular development, RA signalling has many other roles throughout embryonic development^46^. To determine if ID causes a more general upregulation of RA signalling in the developing embryo, we investigated if the development of any other RA-dependent systems were also perturbed. We first investigated the origins of the subcutaneous oedema observed at E15.5. This is a relatively common phenotype in embryonic lethal and sub-viable knockout mouse models. For example, in an unbiased survey of embryonic lethal mouse strains, 24/42 had subcutaneous oedema at E14.5^47^. In humans, this phenotype is called *hydrops fetalis* and can arise from embryonic anaemia or cardiovascular defects, resulting in cardiac failure and a generalised fluid build-up^48^. In our study, we assessed 42 ID E15.5 embryos for both oedema and heart defects. 26 were concordant and 16 discordant, thus there was no correlation between the presence of oedema and heart defects (P<0.0001). Alternatively, oedema can be caused by a failure of normal lymphatic development. This process requires RA signalling, and genetic knockout of *Cyp26b1* results in an identical phenotype to that observed in ID embryos^39^. Therefore, we investigated whether lymphatic development was compromised in ID embryos. The blood and lymphatic vasculature were visualised in E14.5 dorsal back skin using CD31, PROX1 and NRP2 antibodies^49^ (Figure 4a,b). Lymphatic vessels in ID embryos had significantly increased in vessel diameter (Figure 4c) and the vessels had more PROX1-positive nuclei (compare Figure 4 panels a’’ and b’’). The degree of increase in lymphatic vessel diameter (33 -> 52 um) was similar to other mouse models with perturbed lymphatic development^39,50^. By contrast, inter-vessel distance was not significantly altered (Figure 4d), nor were there changes in the patterning of the blood vasculature (Figure 4 a’’’,b’’’). These data suggest that the changes in lymphatic vasculature may be due to a primary defect in lymphatic development, rather than to increased dilation of the vessels subsequent to cardiac failure. In control E11.5 embryos, RA signalling is activated between the dorsal aorta and the ventro-medial side of the cardinal vein (Bowles et al^39^; Figure 4e,f). This is on the opposite wall of the cardinal vein from where lymphatic endothelial cell (LEC) progenitors arise. In ID embryos, RA signalling was increased and expanded dorsally (Figure 4g,h). This is indistinguishable from the phenotypes observed in *Cyp26b1* null embryos, where RA signalling is also increased^39^.

**Figure 4.**
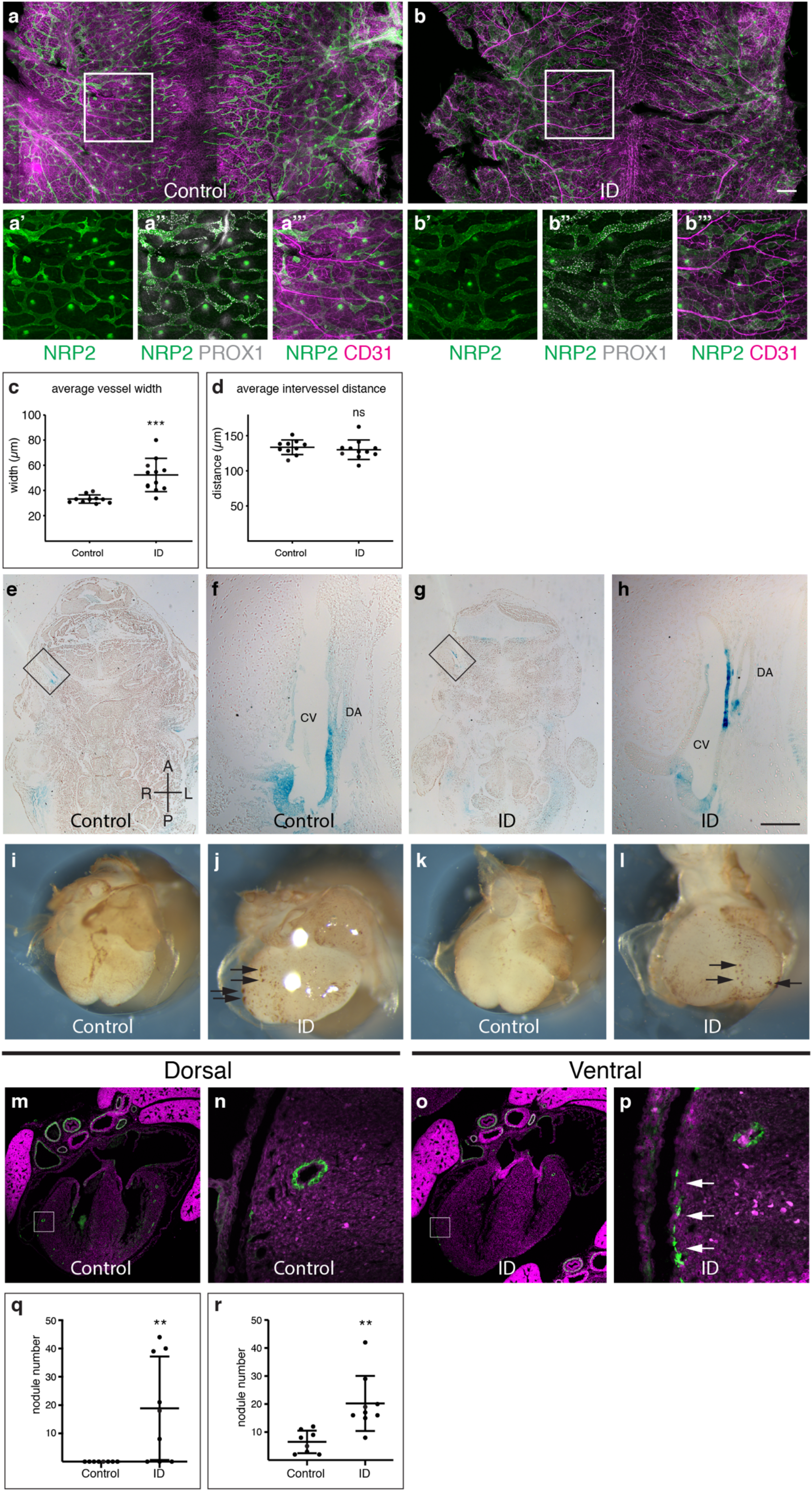
Lymphatic and coronary vasculature development is perturbed in ID embryos. (a-m) Lymphatic development. (a-d) Comparison of NRP2, PROX1 and CD31 expression in back skin from control (a) and ID (b) E14.5 embryos. Magnified views of the boxed areas are shown in panels a’-a’’’ and b’-b’’’, respectively. Quantification of the average lymphatic vessel width (c) and average intervessel distance (d). (e-h) Comparison of RA signalling levels in frontal sections of control (e) and ID (g) E11.5 embryos carrying the RARE-LacZ transgene. (f,h) Magnified views of the boxed areas in e and h, respectively. (i-r) Coronary vasculature development. Comparison of CD31 staining of E14.5 hearts from control (i,k) and ID (j,l). Black arrows indicate endothelial nodules. Quantitation of endothelial nodule number on the dorsal (q) and ventral (r) surfaces. (m-p) Comparison of CD31 (green) expression in the coronary vasculature of control (m) and ID (o) E17.5 embryos. Nuclei were stained with TO-PRO-3 (magenta). Magnified views of the boxed areas are shown in panels n and p, respectively. White arrows indicate ectopic CD31 staining. ns, not significant, ** P<0.01, *** P<0.001. Scale bar = 715 μm (a,b); 370 μm (a’-b’’’); 755 μm (e,g); 95 μm (f,h); 680 μm (i-l); 640 μm (m,o); 64 μm (n,p). CV, cardinal vein; DA, dorsal aorta.

We next examined the development of the coronary vasculature. This develops through a stepwise vasculogenic program. Firstly, an immature vessel plexus forms, and then this is remodelled into a mature vascular bed^51^. The coronary endothelial progenitors arise mostly from the *sinus venosus* (SV) and the endocardium^52^. The epicardium, a layer of cells that migrates over the heart surface from E9.0, is also required for coronary vascular development^53^. It secretes trophic factors, including RA, that stimulate myocardial growth and coronary plexus development. It also contains epicardial-derived progenitor cells (EPDCs) that give rise to cardiac fibroblasts and vascular smooth muscle cells (VSMC) that stabilise the coronary vasculature. EPDCs also contribute to the lateral AV cushions^54^, which were missing in some ID embryos (Figure 1p). 8/8 control hearts at E14.5 had normal coronary plexus formation on both dorsal and ventral surfaces (Figure 4i,k). By contrast, 8/8 ID hearts showed a reduced plexus area, and large numbers of endothelial nodules were present on both heart surfaces (Figure 4j,l, black arrows). This is similar to models of perturbed coronary vascular development due to increased RA signalling^55,56^. By E17.5, 6/6 control E17.5 hearts had clear CD31-positive endothelial tubes throughout the myocardium (Figure 4m,n). By contrast, 6/6 E17.5 ID embryos had fewer obvious endothelial tubes in the myocardium. Instead, patches of CD31-positive cells were visible on the surface of the heart (Figure 4p, white arrows). This phenotype is very similar to that of *Dhrs3* null embryos with increased RA signalling^56^. However, despite the abnormal patterning of distal coronary vessels, the patterning of the proximal coronary vessels was normal. 15/16 E15.5 ID embryos had correctly positioned coronary ostia, although 6/16 had a single additional coronary ostia unilaterally (1 right and 5 left), in line with the previously noted incidence in the C57BL/6J strain^57^. In the absence of normal epicardial formation, myocardial growth is often reduced. Fittingly, ventricular compact myocardium thickness was significantly reduced in E15.5 ID embryos (Extended Figure 2a). Typically, embryos with faulty epicardium and/or coronary vascular development die *in utero* between E12.5-15.5, thus these observations may explain the embryonic lethality in ID embryos.

Finally, somite segmentation is also regulated by RA signalling. Excess RA causes abnormalities in vertebral patterning and delayed ossification^38^. We therefore examined the developing skeletal cartilage in E14.5 embryos by alcian blue staining. 29/33 ID embryos had mild vertebral segmentation defects, including fused lamina, missing pedicles and split vertebral bodies (Extended Figure 4i, 0/21 controls, p<0.0001).

In summary, ID embryos have defects in a variety of tissues that require RA signalling for normal patterning, and these defects are similar to those in genetic models of increased RA signalling. Thus, our data supports the hypothesis that ID results in a wide-spread disruption of RA signalling in the developing embryo.

## Dietary supplementation mid-gestation rescues the heart and lymphatic phenotypes

Clinically, ID can be treated rapidly and effectively by intravenous administration of ferric carboxymaltose or dietary supplementation. Therefore, it would be useful to know if, and when, during pregnancy that iron supplementation might rescue embryonic defects. We transferred pregnant ID mice from low iron to normal diet (200 ppm iron) 7-9 days post-mating. In each case, maternal haemoglobin levels returned to normal by E15.5 (Extended Figure 5a). Diet change on days 8 or 9 had no significant effect on the prevalence of either oedema or embryonic lethality at E15.5 (Figure 5a). By contrast, diet change on day 7 resulted in a significant reduction in both oedema and embryonic lethality (2/53 embryos abnormal or dead, compared to 38/80 ID embryos, P<0.0001; and 0/58 controls, P=0.2257). Furthermore, 0/26 of viable E7.5-rescued embryos examined at E15.5 had heart defects (30/42 ID embryos, P<0.0001; and 1/37 controls, P=0.5873; Extended Table 1), indicating a complete rescue of the heart phenotype in these embryos. Iron uptake from the gut is swift, taking only a few hours^58^, and is even faster in ID animals with low hepcidin levels^59^. Once in the maternal bloodstream, iron is transferred to the embryo in <6 hours^60^. Thus, phenotypic rescue by returning mothers to the iron-replete diet on E7.5, but not later, suggests that the critical period of embryonic development that is sensitive to ID is approximately E8.5. This is identical to studies of RA exposure, where embryonic heart development is most vulnerable at E8.5^61–63^. It is also broadly similar to our previous studies of embryonic hypoxia, which showed a peak in vulnerability of SHF cells to hypoxia at E9.5^5^. This stage of development is when SHF cells migrate into the OFT and when pharyngeal arch morphogenesis takes place. It also corresponds to the initial movement of epicardial cells to the heart’s surface^64^ and the induction of lymphatic endothelial cells in the wall of the cardinal veins^65^. Thus, restoration of normal RA signalling at this stage is likely to explain the rescued cardiac, lymphatic and coronary vessel defects.

**Figure 5.**
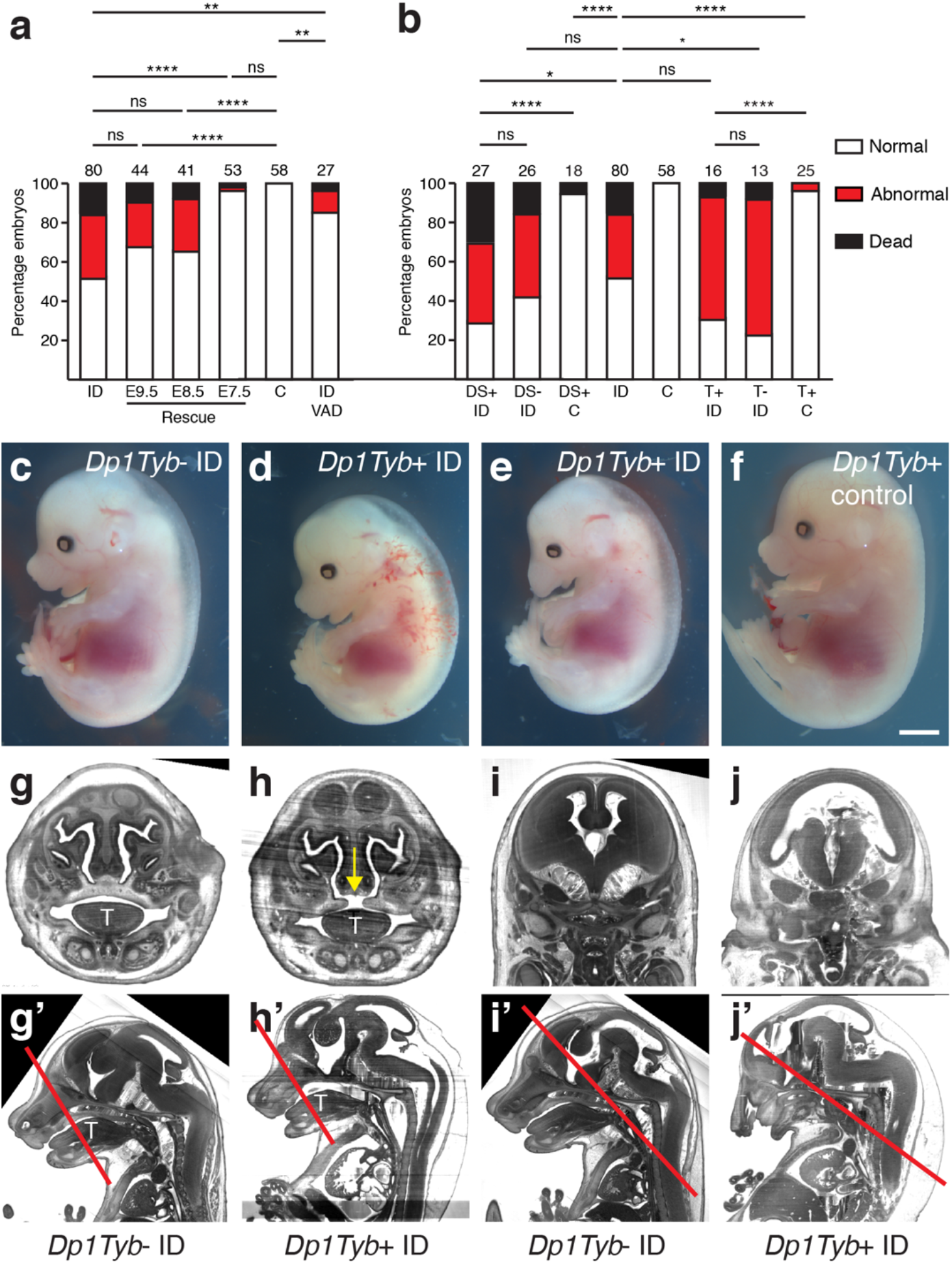
Phenotypic rescue and gene-environment interaction. (a) Phenotypic rescue. Pregnant mice were returned to iron-replete diet between E7.5-E9.5 or fed continuously on an iron and vitamin A deficient diet from weaning and throughout pregnancy (ID/VAD). Histograms showing the percentage of embryos that were dead, abnormal or normal at E15.5. (b) Investigation of gene-environment interaction. Iron deficient (ID) and control (C) C57BL/6J females were crossed with males carrying the *Tbx1*^*null*^ allele (T), or the *Dp1Tyb* duplication (DS). Histograms showing the percentage of embryos that were dead, abnormal or normal at E15.5. (c-f) Representative images of *Dp1Tyb*^*+*^ control (c), *Dp1Tyb* control ID (d), *Dp1Tyb*^*+*^ ID with holoprosencephaly (e), *Dp1Tyb+* control (f), E15.5 embryos. (g-j) *Dp1Tyb*+ embryos from ID mothers have craniofacial defects. Representative 3D reconstructed frontal sections of HREM data from *Dp1Tyb*− ID (g,i) and *Dp1Tyb*+ ID (h,j) E15.5 embryos showing examples of failed secondary palate fusion (h, yellow arrow) and holoprosencephaly (j). (g’-j’) Sagittal sections from the same embryos showing the location of the sections in g-j (red lines). note that panels g and i are the same control embryo. T, tongue. ns, not significant, ** P<0.01, *** P<0.001, **** P<0.0001. Scale bar = 2 mm (c-f); 700 μm (g,h); 1.2 mm (I,j); 1.6 mm (g’-I’).

## Mothers deficient for both iron and vitamin A produce normal embryos

We next tested our mechanistic hypothesis by further altering the maternal diet. RA is produced from dietary vitamin A. It has been known since the 1930s that maternal vitamin A deficiency (VAD) disrupts development^66^, and this is now known to be caused by reduced embryonic RA signalling. We reasoned that embryos of a mother fed a diet low in iron (leading to increased RA signalling) and deficient in vitamin A (leading to reduced RA signalling) might result in relatively normal levels of RA, and thus restore normal embryonic development. Female C7BL/6J strain mice were weaned onto, and maintained continuously on, a diet containing low iron (2-6 ppm) and no added vitamin A (control and ID diets both have 15 IU/g vitamin A). As before, these mice had significantly reduced blood haemoglobin, confirming that VAD did not affect iron metabolism (Extended Figure 5a). These mice were mated to control C57BL/6J males, then maintained on the ID/VAD diet until embryo collection on E15.5. μMRI imaging confirmed that embryos had reduced liver iron levels (Extended Figure 5c). Strikingly, only 1/27 embryos were dead and 3/27 had oedema (Figure 5a). This is a significantly lower rate of death and abnormality than ID embryos (38/80 ID embryos, P=0.0019), although not a complete phenotypic rescue (0/58 controls, P=0.0087). Furthermore, only 3/25 of these embryos had heart defects, compared to 30/42 ID (P<0.0001) and 1/37 control embryos (P=0.1752), indicating an almost complete rescue of heart defects. These data provide further evidence that the defects observed in embryos from ID embryos are due to increased RA signalling.

## Gene-environment interactions increase the penetrance and expressivity of embryonic defects

In human CHD, even in families with a known monogenic cause, there is often an array of different cardiac defects between individuals with the same causative mutation (variable expressivity), while others with the same mutation do not develop CHD at all (variable penetrance)^67^. This suggests that CHD phenotypes are commonly affected by genetic or environmental modifiers. Previously we have shown in mouse that short-term gestational hypoxia is one such modifier, increasing the prevalence and severity of heart defects in genetically-susceptible embryos, as well as causing heart failure and embryonic lethality^68,69^. We hypothesised that ID might cause a similar gene-environment interaction (GxE). Two of the most common human genetic syndromes that include CHD are 22q11.2 deletion syndrome^70^ and Down syndrome (DS)^71^. In both cases, the types of heart defects presented clinically are similar to those of the ID model. Furthermore, in *Tbx1* null embryos all three *Cyp26* genes are downregulated, suggesting that RA signalling may be altered in 22q11.2 deletion syndrome^40^. It has long been appreciated that there is considerable variation in the severity and penetrance of cardiovascular phenotypes in human patients with either syndrome. To test the hypothesis that this variation might be controlled by an environmental factor such as ID, we crossed male C57BL/6J background mice heterozygous for either a *Tbx1* null allele^72^ (a model of 22q11.2 deletion syndrome), or the *Dp1Tyb* duplication^73^ (a model of DS), with ID females and analysed embryonic phenotypes at E15.5. In both cases, we confirmed that maternal haemoglobin levels were significantly reduced (Extended Figure 5a). There was no significant increase in the prevalence of embryonic death and oedema, heart defects or aortic arch abnormalities in *Tbx1*^*+/null*^ ID embryos (Figure 5b; Extended Table 1; Extended Table 2). Thus, maternal low iron status is not likely to be a risk factor for increasing penetrance or severity of cardiovascular phenotypes in 22q11.2 deletion syndrome. Similarly, E15.5 ID embryos carrying the *Dp1Tyb* duplication (*Dp1Tyb*+ ID) had the same prevalence of death and abnormality as *Dp1Tyb−* ID control littermates (Figure 5b, 19/27 *Dp1Tyb+* ID, 15/26 *Dp1Tyb−* ID littermates, P=0.2498). Strikingly however, affected embryos had more severe subcutaneous oedema, and their lymphatics were more frequently blood-filled (Figure 5d; 10/11 *Dp1Tyb+* ID oedemic embryos with blood-filled lymphatics compared to 4/11 *Dp1Tyb*− ID, P=0.0119). In addition, we observed craniofacial defects in surviving *Dp1Tyb*+ ID embryos. 10/19 of these embryos had a failure of secondary palate fusion (1/39 wild type ID controls, P<0.0001, Figure 5h), and a further 2 embryos had holoprosencephaly (Figure 5e,j). By contrast, only 1/18 *Dp1Tyb*+ control embryos was dead at E15.5, and none of the survivors had oedema, holoprosencephaly or a failure of secondary palate fusion (Figure 5f). This induction of novel phenotypes is an indication that we have identified a *bona fide* gene-environment interaction, rather than a simple additive effect. We also assessed heart morphology in surviving embryos at E15.5. 16/18 ID *Dp1Tyb*+ ID embryos had heart defects (Extended Table 1), compared to 10/19 *Dp1Tyb*− ID littermates (P=0.0186) and 1/16 *Dp1Tyb*+ controls (P<0.0001). Furthermore, *Dp1Tyb*+ ID embryos had significantly more AVSDs (10/18) than *Dp1Tyb*− ID embryos (3/19, P=0.0135), *Dp1Tyb+* control embryos (0/16, P=0.0003) or ID embryos (5/42, P=0.0008). Finally, 4/18 Dp1Tyb+ ID embryos had type I PTA, which was never observed in *Dp1Tyb*− ID control, *Dp1Tyb+* control or wild type ID control embryos. Thus, we conclude that maternal ID leads to an increase in the penetrance and severity of heart defects in *Dp1Tyb*+ mouse embryos. By contrast, there was no significant change in the prevalence of AA anomalies between *Dp1Tyb*+ ID and *Dp1Tyb*− ID embryos (Extended Table 2). In conclusion, ID may be a significant modifier of heart, lymphatic and/or craniofacial phenotype in children with DS.

## Discussion

Here, we have identified a completely new environmental teratogen: maternal ID anaemia. Clinically, this is potentially of great importance, since iron deficiency is the most common micronutrient deficiency worldwide. We have shown in mice that maternal ID causes severe embryonic cardiovascular defects via premature differentiation of a subset of cardiac progenitor cells. At the molecular level, this most likely results from increased retinoic acid signalling. Furthermore, we show that the defects can be rescued by iron administration early in pregnancy, or by reducing vitamin A intake in iron deficient mothers. Although our results do not formally distinguish between ID or generalised anaemia as the cause of the defects, we believe it is more likely to be ID. The evidence for this is two-fold. Firstly, we have demonstrated that maternal ID phenocopies embryonic loss of CYP26 activity in the heart, coronary vessels and lymphatic system, phenotypes that in each case result from increased embryonic RA signalling. CYP26 is an iron-dependent enzyme, therefore ID may well partially reduce its enzymatic activity, resulting in mildly increased RA signalling. Secondly, human epidemiological studies suggest that maternal anaemia only increases offspring CHD risk minimally (adjusted odds ratio (OR) 1.2^74–76^), whereas low iron intake in the first trimester (with or without overt anaemia) has an adjusted OR of offspring CHD of up to 5.0^10^. Our hypothesis that ID causes mildly increased RA signalling is supported by studies of the effects of exposure of pregnant mice to excess RA^61–63,77^. Here, administration of low doses of RA results in the same phenotypes as ID at E15.5, including isolated membranous VSD and DORV. By contrast, administration of high doses of RA causes TGA, which we did not observe in ID embryos. Furthermore, low doses of RA also cause highly similar morphological defects to ID earlier in development: shortening and rotational defects of the OFT at E10.5; hypoplasia and dysplasia of the proximal OFT cushions (but not the AVC cushions) and hypoplastic or absent aortic ICC at E12.5^61–63^; and thin ventricular myocardium at E13.5^77^. However, one feature of our hypothesis that is difficult to explain is why ID has a relatively minor effect on embryogenesis. Iron is required for the function of almost 400 human proteins^78^, and one might imagine that the activities of many of these proteins would also be affected in ID embryos. Why the CYP26 enzymes might be particularly sensitive to reduced iron levels in the embryo remains unclear.

Our observations have important clinical implications. We induced maternal ID via environmental modification. However, maternal ID can also arise from genetic insufficiency and this could potentially have similar effects on embryonic development. The induction of identical phenotypes in different individuals by genetic, environmental or gene-environment interaction is called phenocopying. This has previously been suggested to occur in some types of human congenital abnormalities, for example congenital NAD deficiency disorder^79,80^. Our results suggest that some cases of CHD or craniofacial defects might arise from maternal mutations in the almost 40 genes required for iron transport and/or metabolism. To address how common such mutations might be in humans, we examined the Genome Aggregation Database^81^. This is an aggregate of human exome and genome sequencing data from 141,456 individuals without severe paediatric diseases. We identified 771 predicted loss-of-function variants in these genes. Individuals carrying these variants might be predisposed to developing iron deficiency, and thus may have increased risk of having offspring with birth defects.

Our discoveries may also explain some of the variable penetrance of CHD and cleft palate in children with DS. Our unexpected observation of a strong gene-environment interaction resulting in a failure of secondary palate fusion supports our hypothesis that ID causes increased RA signalling. *Cyp26b1* null mouse embryos have fully-penetrant cleft palate^82,83^, and high maternal doses of Vitamin A can also induce cleft palate in mouse^84^. In addition, people with DS have an increased risk of cleft palate^85^. Thus the combination of mildly increased RA due to ID, combined with a genetic susceptibility due to DS, may explain our results. This hypothesis is further supported by the presence of PTA in some *Dp1Tyb+* ID embryos. This phenotype is typical of offspring of mothers administered high doses of RA^61–63^, but was not observed in ID alone or *Dp1Tyb+* control embryos. However, the link between *Dp1Tyb* and RA signalling is not obvious. The duplicated region contains 172 protein coding genes, of which at least three have been associated with RA signalling: *Nrip1*, *Runx1* and *Ripply3*. Intriguingly, Ripply3 is a transcriptional co-repressor of Tbx1^86^ and is a direct target of RA signalling^87^. Thus, in *Dp1Tyb+* ID embryos, slightly increased RA signalling coupled with an extra copy of *Ripply3*, may cause repression of Tbx1 target genes, resulting in a phenocopy of *Tbx1* null phenotypes including cleft palate^88^ and more severe cardiovascular defects. Finally, the clinical relevance of our gene-environment interaction observations could be relatively easily tested by a prospective clinical study examining the clinical effects of maternal peri-conception iron status on the phenotypes of children with DS.

If maternal ID does indeed lead to increased RA signalling in the embryo, then ID may have a particularly great impact on women with severe cystic acne. Isotretinoin (Roaccutane®) is a common and effective treatment for this condition, but it is well-known that this isoform of RA is highly teratogenic^42^. Our results suggest that ID may further exacerbate the teratogenicity of isotretinoin. Best practice indicates use of two reliable methods of contraception for one month before starting treatment, and for one month after treatment has stopped. However, there is evidence of a significant non-compliance rate in these patients^89^, and thus it might be of benefit for patient iron status to be monitored during treatment to reduce the potential effects of excess RA on an unintended pregnancy.

Our finding that combined iron and vitamin A deficiency substantially rescues the cardiovascular defects may explain the disparity between animal experiments and epidemiological studies of VAD. There is very strong evidence in animal models that VAD alone is highly teratogenic, but epidemiological studies to date have not found a particularly strong association between VAD and CHD. In the developing world, 15% of pregnant women have VAD^90^ and these women also commonly have ID as well. Our results might suggest that the combination of iron and vitamin A deficiency will balance RA levels, and thus mask any effect of VAD alone on CHD prevalence.

Studies of genetic causes of birth defects are useful on a case-by-case basis to provide information on prognosis, treatment options and recurrence rate. However, understanding particular genetic causes of birth defects has limited value in reducing their overall birth prevalence. By contrast, our elucidation of ID as a potential new environmental risk and/or modifying factor for CHD may guide changes in clinical advice. Current WHO and NICE guidelines have conflicting advice on iron supplementation during pregnancy, with the WHO recommending daily iron supplementation to all pregnant women, whereas NICE recommends supplementation only in cases of substantial ID^91,92^. Our results suggest that all women of child-bearing age should be advised to maintain optimal iron levels. This conclusion is supported by a recent study suggesting that low iron intake during early pregnancy in humans increases the risk of offspring CHD by up to 5-fold^10^. Iron supplementation during pregnancy is unlikely to have a deleterious effect on embryonic development. Pregnant mice loaded with excess iron have increased levels of the hormone hepcidin. Hepcidin prevents the release of excess iron into the maternal circulation, and these increased levels protect her embryos from iron overload^60^. Thus, recommending iron administration as soon as pregnancy is suspected may be a safe and effective strategy for reducing risk of CHD worldwide.

---

**Extended Figure 1.**
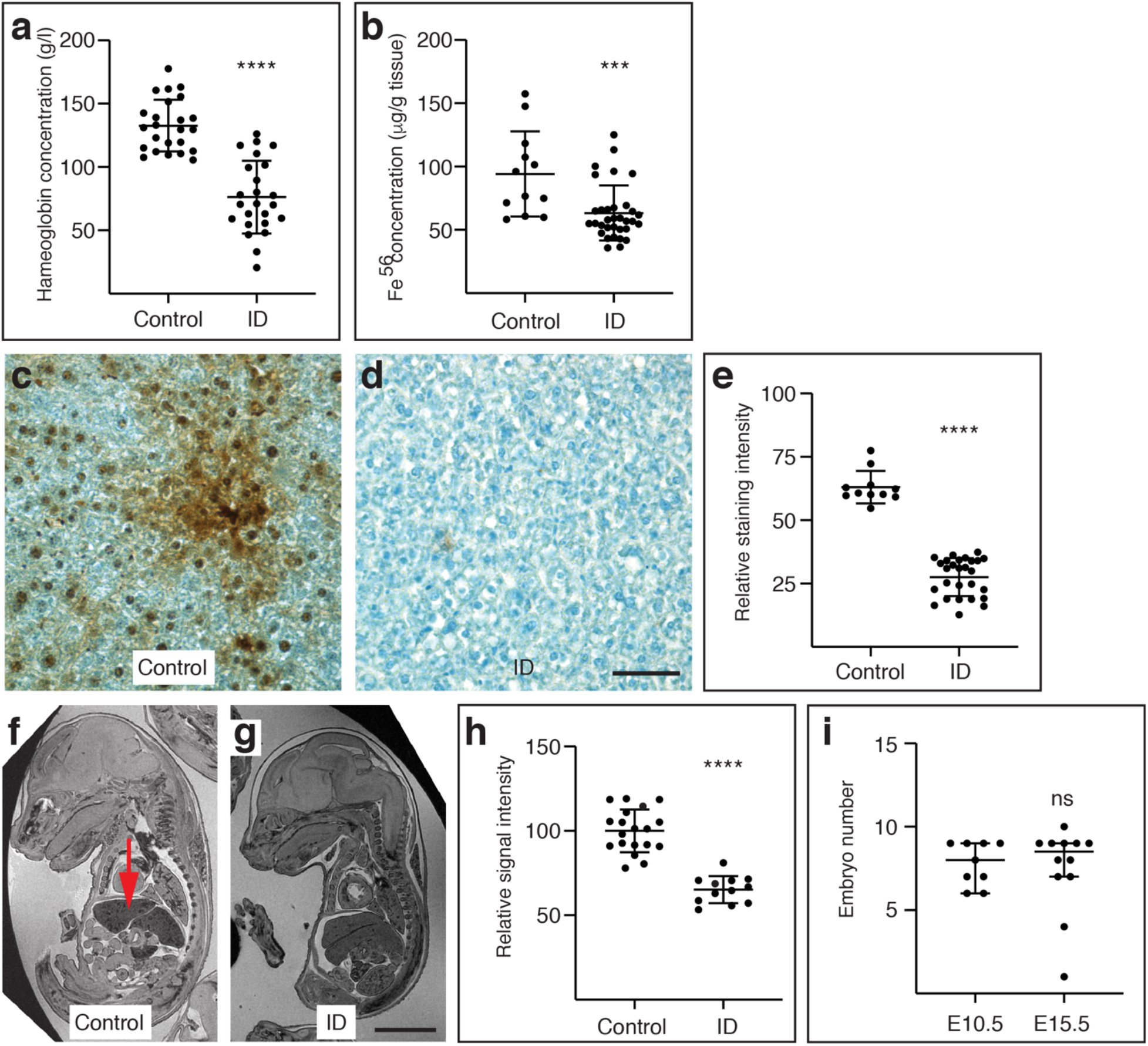
Mice and embryos continuously fed a low iron diet from weaning are ID and anaemic. (a) Comparison of blood haemoglobin concentrations between adult female mice fed since weaning on control (200 ppm iron) or low iron (2-6 ppm iron) diet. (b) Comparison of Fe^56^ concentration in liver samples measured by Inductively Coupled Plasma Mass Spectrometry. (c-e) Representative images showing relative DAB-enhanced Perls staining of liver sections from control (c) or ID (d) mice. (e) Quantitation of Perls staining. (f-h) Representative images showing relative liver MRI contrast from control (f) or ID (g) E15.5 embryos. The embryonic liver is shown by a red arrow. (g) Quantitation of liver MRI contrast. (i) Comparison of litter size (including dead embryos) in ID mothers between E10.5 and E15.5. ns, not significant, *** P<0.001, **** P<0.0001. Scale bars are 50 μm (c,d) and 3 mm (f,g).

**Extended Figure 2.**
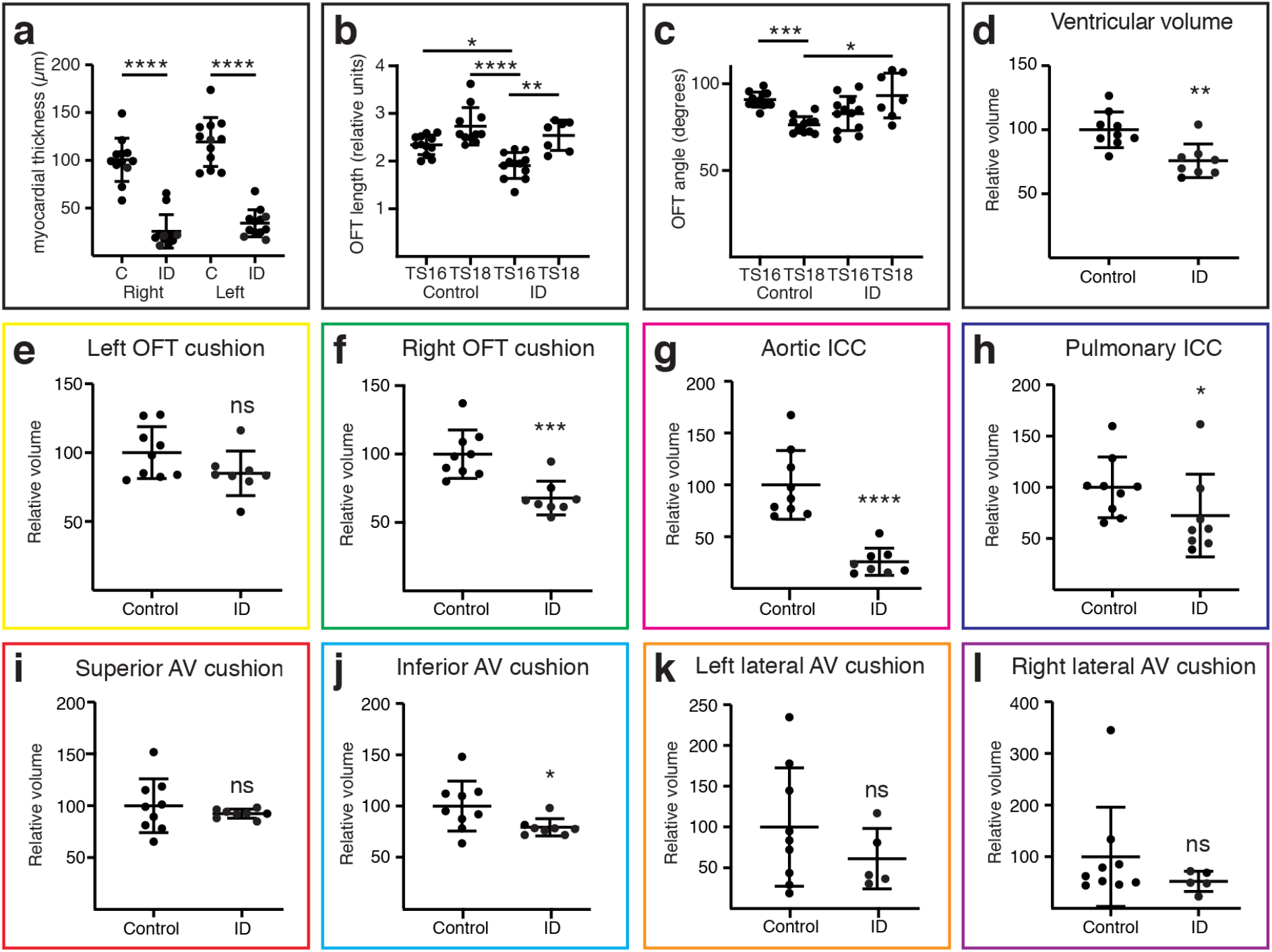
Maternal ID causes cardiovascular defects. (a) Quantification of the ventricular myocardial thickness of hearts from E15.5 control and ID embryos. (b-c) Quantification of length (b) and angle between distal and proximal portions of the OFT (c) from control and ID E10.5 embryos. (d-l) Quantitation of whole ventricle and cardiac cushion volumes from 3D Amira models of manually-segmented HREM data from E12.5 embryos. The panel border colour corresponds to the cushion colour in Figure 1 (panels h-q).* P<0.05, ** P<0.01, *** P<0.001, **** P<0.0001.

**Extended Figure 3.**
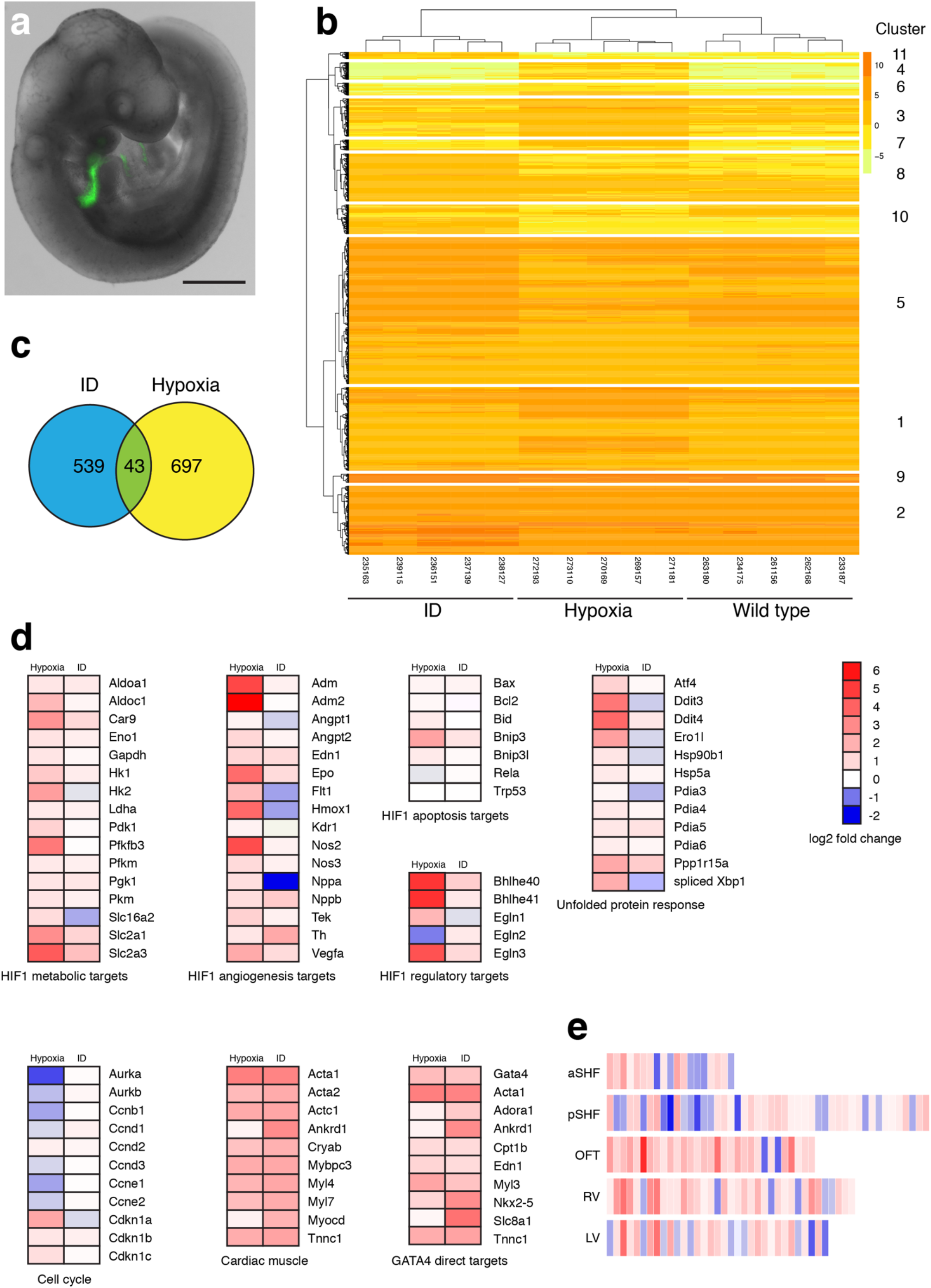
Transcriptomic analysis of anterior SHF cells. (a) Representative E9.5 embryo showing the expression domain of the Mef2c-AHF-GFP transgene. Scale bar is 500 μm. (b) Unsupervised hierarchical clustering using normalised gene expression levels ordered samples into the correct condition groups. Only differentially expressed genes are shown with adjusted p value <0.01, B >1 and differentially expressed <> 2-fold. (c) Comparison of genes differentially expressed relative to control samples shows little overlap between ID and hypoxia samples. Analysis was restricted to genes with adjusted p value <0.01, B >1 and differentially expressed <> 2-fold. (d) Comparison of expression changes relative to control samples of ID and hypoxia samples. Heat maps show the average fold-change between ID or hypoxia and control samples of selected genes involved in the hypoxia response, the unfolded protein response, cell cycle control, cardiac muscle proteins and GATA4 direct targets. (e) The ID transcriptome is more closely related to the OFT cluster than the aSHF cluster from single-cell RNA-Seq of heart and cardiac progenitor cells at E9.25 (de Soysa et al^33^). Heat maps show every gene in each de Soysa cluster with significant (p<0.05) fold-change between ID and control samples. Statistical significance was tested using a hypergeometric test (assuming 20,000 genes in the transcriptome) with Bonferroni correction to control the familywise error rate. P values: OFT 3.58×10^−6^; RV 3.29×10^−4^; aSHF 3.54×10^−4^; pSHF 1.87×10^−3^; LV 1.05×10^−2^). aSHF, anterior second heart field; pSHF, posterior second heart field; OFT, outflow tract; RV, right ventricle; LV, left ventricle.

**Extended Figure 4.**
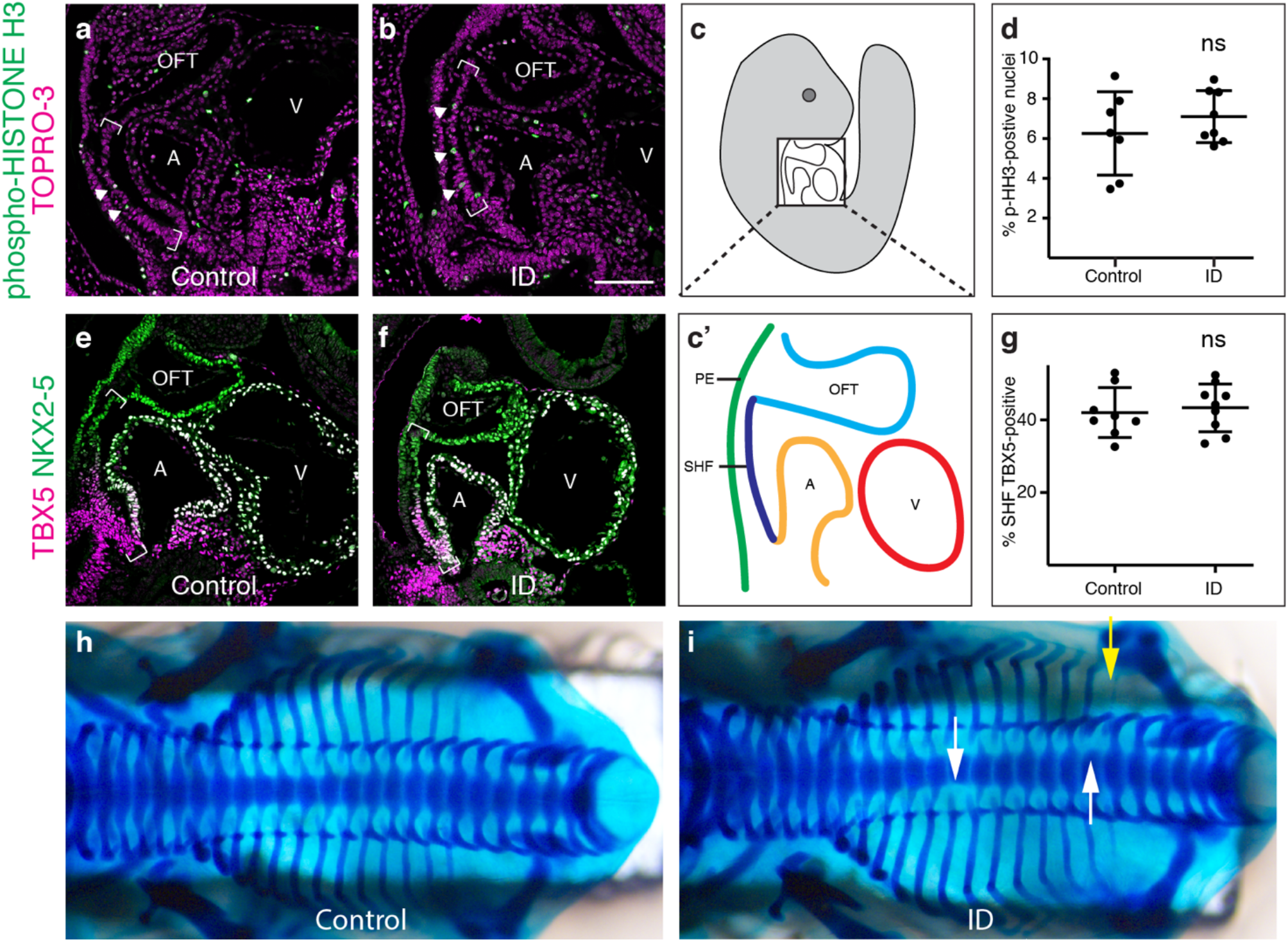
Effects of ID on cardiac progenitor proliferation, TBX5 expression and skeletal patterning. (a-d) Comparison of expression levels of phospho-Histone H3 (pHH3, green) in sagittal sections of control (a) and ID (b) E9.5 mouse embryos by immunohistochemistry. Nuclei were stained with TO-PRO-3 (magenta). Location of the SHF is indicated by brackets. (c,c’) Diagrams indicating the relative positions of the SHF (dark blue), pharyngeal endoderm (green), OFT (light blue) and left ventricle (V, red) and left atrium (A, orange) in a sagittal section of an E9.5 embryo. (d) Quantification of number of pHH3-positive SHF cells. (e-g) Comparison of expression levels of TBX5 (magenta) and NKX2-5 (green) in control (e) and ID (f) E9.5 mouse embryos by immunohistochemistry. Location of the SHF is indicated by brackets. (g) Quantification of the percentage of TBX5-positive SHF. (h,i) Alcian blue staining of cartilage in control (h) and ID (i) E14.5 embryos. Missing pedicles (black arrows) and rib (yellow arrow) are indicated. ns = not significant. Scale bar = 130 μm (a,b,e,f), 520 μm (h,i).

**Extended Figure 5.**
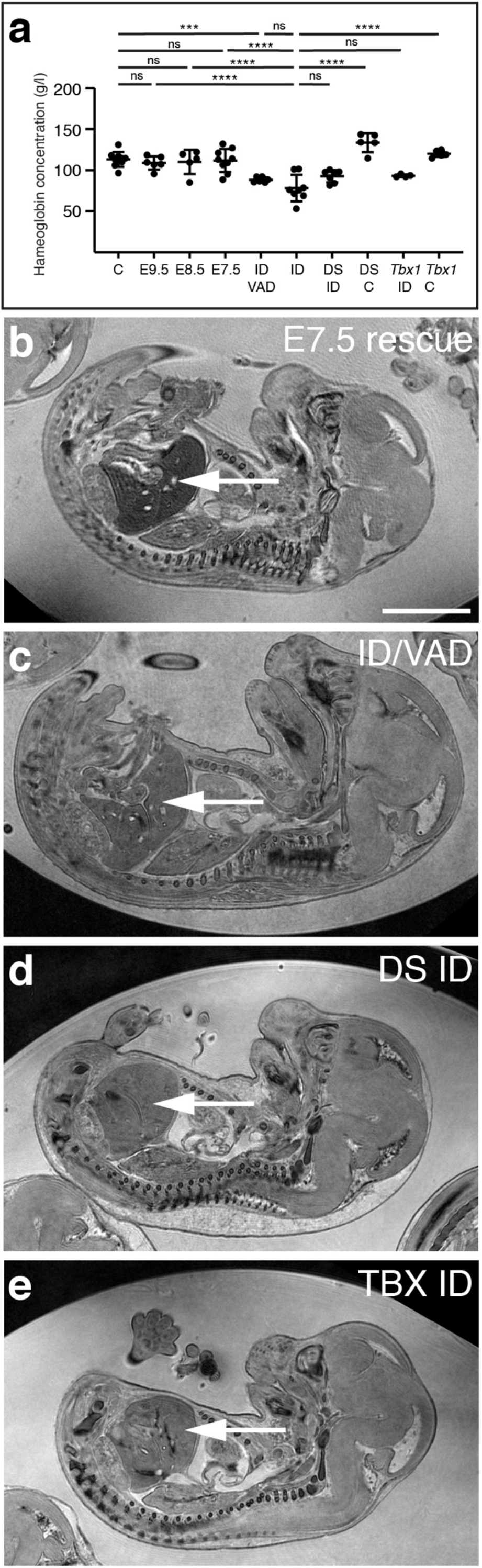
Effects of maternal diet on blood haemoglobin levels and embryonic liver iron. (a) Blood haemoglobin concentrations in pregnant adult C57BL/6J strain female mice measured at E15.5. C, mice fed control diet; E9.5, E8.5, E7.5, mice fed from weaning on a low iron diet, then returned to control diet at the indicated stage of pregnancy; ID/VAD, mice fed from weaning on a low iron and Vitamin A-deficient diet; ID, mice fed from weaning on a low iron diet. DS = C57BL/6J females crossed to *Dp1Tyb*+ males; Tbx1 = C57BL/6J females crossed to *Tbx1*^*+/null*^ males. (b-e) Representative images showing relative liver MRI contrast in E15.5 embryos from (b) ID mothers returned to control diet on E7.5; (c) mothers fed a low iron and vitamin A-deficient diet; (d) *Dp1Tyb*+ embryos from ID mothers; (e) *Tbx1*^*+/null*^ embryos from an ID mother. The embryonic livers are shown by a white arrow. Scale bar = 3 mm.

**Extended Table 1.** Summary of types of heart defects observed in E15.5 embryos. ASD, atrial septal defect; mVSD: membranous ventricular septal defect; musVSD: muscular ventricular septal defect; AVSD, atrioventricular septal defect; OFT, outflow tract defects (includes overriding aorta and double-outlet right ventricle); PTA, persistent *truncus ateriosus*. † one with type I PTA; ‡ three with type I PTA

| Types of defects |  | Control | ID | E7.5 rescue | ID/VAD |  | *Dp1Tyb*<sup>+</sup> ID | *Dp1Tyb*<sup>-</sup> ID | *Dp1Tyb*<sup>+</sup> control |  | *Tbx1*<sup>+/*null*</sup> ID | *Tbx1*<sup>+/*+*</sup> ID | *Tbx1*<sup>+/*null*</sup> control |
| --- | --- | --- | --- | --- | --- | --- | --- | --- | --- | --- | --- | --- | --- |
| Single | ASD | 0 | 0 | 0 | 0 |  | 0 | 0 | 0 |  | 0 | 0 | 0 |
|  | mVSD | 1 | 5 | 0 | 0 |  | 0 | 1 | 0 |  | 1 | 1 | 0 |
|  | muscVSD | 0 | 11 | 0 | 0 |  | 3 | 1 | 1 |  | 4 | 3 | 0 |
|  | AVSD | 0 | 0 | 0 | 0 |  | 1 | 0 | 0 |  | 0 | 0 | 0 |
| Multiple | OFT + mVSD | 0 | 1 | 0 | 2 |  | 0 | 2 | 0 |  | 1 | 0 | 0 |
|  | mVSD + muscVSD | 0 | 4 | 0 | 0 |  | 0 | 1 | 0 |  | 2 | 1 | 0 |
|  | OFT + mVSD + muscVSD | 0 | 4 | 0 | 0 |  | 3† | 2 | 0 |  | 4 | 1 | 1 |
|  | mVSD + AVSD | 0 | 0 | 0 | 0 |  | 0 | 0 | 0 |  | 0 | 0 | 0 |
|  | OFT + mVSD + AVSD | 0 | 0 | 0 | 0 |  | 1 | 1 | 0 |  | 0 | 0 | 0 |
|  | muscVSD + AVSD | 0 | 4 | 0 | 0 |  | 0 | 0 | 0 |  | 0 | 1 | 0 |
|  | OFT + muscVSD + AVSD | 0 | 1 | 0 | 1 |  | 8‡ | 2 | 0 |  | 1 | 2 | 0 |
| Total abnormal |  | 1 | 30 | 0 | 3 |  | 16 | 10 | 1 |  | 13 | 9 | 1 |
| Total normal |  | 36 | 12 | 26 | 22 |  | 2 | 9 | 15 |  | 3 | 3 | 18 |
| P value vs control |  | - | <0.0001 | 0.5873 | 0.1752 | P value vs *Dp1Tyb*<sup>+</sup> control | <0.0001 | 0.0037 | - | P value vs *Tbx1*<sup>+/*null*</sup> control | <0.0001 | <0.0001 | - |
| P value vs ID |  | - | - | <0.0001 | <0.0001 | P value vs *Dp1Tyb*- ID | 0.0186 | - | - | P value vs *Tbx1*<sup>+/+</sup> ID | 0.5208 | - | - |

**Extended Table 2.** Summary of aortic arch morphology at E15.5.

| Types of defects | control | ID | E7.5 rescue | ID/VAD |  | *Dp1Tyb+* ID | *Dp1Tyb-* ID | *Dp1Tyb+* control |  | *Tbx1*<sup>+/*null*</sup> ID | *Tbx1*<sup>+/*+*</sup> ID | *Tbx1*<sup>+/*null*</sup> control |
| --- | --- | --- | --- | --- | --- | --- | --- | --- | --- | --- | --- | --- |
| IAA | 0 | 3 | 0 | 0 |  | 4 | 1 | 0 |  | 0 | 0 | 1 |
| A-RSA | 0 | 1 | 1 | 1 |  | 1 | 3 | 0 |  | 4 | 1 | 4 |
| R-LCC | 0 | 1 | 0 | 0 |  | 2 | 0 | 1 |  | 0 | 0 | 0 |
| right-sided AA | 1 | 1 | 0 | 0 |  | 2 | 0 | 1 |  | 2 | 0 | 1 |
| abnormal | 1 | 5 | 1 | 1 |  | 7 | 4 | 1 |  | 6 | 1 | 6 |
| normal | 36 | 35 | 24 | 24 |  | 11 | 15 | 14 |  | 10 | 11 | 13 |
| P value vs control | - | 0.1189 | 0.6478 | 0.6478 | P value vs *Dp1Tyb+* control | 0.0375 | 0.2507 | - | P value vs *Tbx1*<sup>+/*null*</sup> control | 0.4946 | 0.1430 | - |
| P value vs ID | - | - | 0.2456 | 0.2456 | P value vs *Dp1Tyb-* ID | 0.2046 | - | - | P value vs *Tbx1*<sup>+/*+*</sup> ID | 0.0908 | - | - |

IAA, interrupted aortic arch; A-RSA, aberrant right subclavian artery; R-LCC, retroesophageal left subclavian artery.
Note that an embryo may have more than one phenotype.

## Methods

### Animals

All animal experiments were compliant with the UK Animals (Scientific Procedures) Act 1986 and approved by the University of Oxford animal welfare review board and the Home Office (project license PB01E1FB3). Mice were housed in an SPF facility free from the major rodent pathogens except *Helicobacter hepaticus*, with a 12:12 hour light/dark cycle, at 19-23 °C, 55 ±10 % humidity, in individually-ventilated cages (Tecniplast UK Ltd, Rushden, UK) containing Grade 4 Aspen Chip bedding (Datesand Ltd, Manchester, UK), cardboard tunnels and Sizzle Pet nesting material (LBS Biotechnology, Horley, UK), with free access to food and tap water. Bedding was changed fortnightly, and animals were assessed daily for welfare. Mice were fed with Teklad 2916 or TD.08713 (control), TD.99397 (iron deficient) or TD.190023 (iron and vitamin A deficient), all from Envigo, Belton, UK. C57BL/6J mice were purchased from Charles River UK. Genetically-modified mouse strains were; *Tg(Mef2c-EGFP)#Krc* (Mef2c-AHF-GFP)^32^; *Tg(RARE-Hspa1b/lacZ)12Jrt* (RARE-LacZ)^45^; *Dp(16Lipi-Zbtb21)1TybEmcf* (*Dp1Tyb*)^73^; and *Tbx1*^*tm1Bld*^ (*Tbx1*^*LacZ*^)^72^. All genetically-modified mice were from colonies that had been backcrossed for more than 10 generations onto the C57BL/6J background, with the exception of the RARE-LacZ strain, which was maintained on a CD-1 background.

### Blood haemoglobin and liver iron content measurement

Blood haemoglobin levels were measured in fresh blood using a HemoCue^®^ Hb 201^+^ according to the manufacturer’s instructions. Values of two separate samples were averaged. Two liver samples were taken per animal at sacrifice and one snap-frozen in liquid nitrogen, and the other fixed overnight in Formalin. Frozen samples were analysed by inductively coupled plasma mass spectrometry (ICP-MS) as previously described^93^. Fixed samples were paraffin embedded, sectioned, and stained by the DAB-enhanced Perls method as previously described^93^. Slides were imaged with a Nikon COOLSCOPE slide scanner and staining intensity was measured by colour deconvolution with the H DAB vector in FIJI 2.0.0-rc-69/1.52p software.

### Embryo and heart morphology assessment

Micromagnetic resonance imaging (μMRI) was performed as previously described^94,95^, using either a Varian 9.4 T VNMRS 20 cm horizontal-bore system (Varian Inc. Palo Alto, CA, USA) or a 11.7 T (500 MHz) vertical magnet (Magnex Scientific, Oxon, UK), both running a Varian/Agilent DDR2 console. Fixed embryos were incubated in 2 mM Magnevist® (Bayer) and embedded in agarose in a Wilmad LabGlass 28-PP-9” tube. Parameters for the 9.4T system were: TR 28 ms, TE 16ms, flip angle 52°, 5 averages, 27×27×27mm^3^ with a matrix size of 512^3^, giving an isotropic resolution of 52×52×52 μm^3^ per voxel. A hard RF pulse (duration 100ms) was used for excitation and receiver bandwidth of 66 kHz. Parameters for the 11.7T system were: TR 16.8 ms; TE 4.5 ms; 50° flip angle (BIR-4 adiabatic); 9 averages, 24×24×32 mm^3^ FOV; 1024×1024×1366 matrix size, 50 kHz bandwidth, 75% partial fourier scheme^96^; and reconstructed with the Berkley advanced reconstruction toolkit (https://mrirecon.github.io/bart/) as 1536×1536×2049 (complex double) voxels with an isotropic 15.6 μm^3^ resolution. In all cases, μMRI was followed by paraffin embedding, sectioning and H&E staining. High resolution episcopic microscopy (HREM) was performed as previously described^97^ using an JB-4 embedding kit (00226-1, Polysciences GMBH, Germany). μMRI and HREM data were analysed using OsiriX MD DICOM viewer version 9.0.2 (Pixmeo), Horos 3.3.6 (https://horosproject.org) and Amira for Life & Biomedical Sciences version 2019.4 (Thermo Fisher Scientific).

### RNA-Seq

1,110-4,566 GFP-expressing viable cells were sorted from E9.5 embryos carrying the Mef2c-AHF-GFP allele^30^ by FACS using a Beckman Coulter MoFlo AstriosEQ with Summit 6.2.7.16492 software. GFP was detected with the 488nm laser and a 513/26 band pass filter, and DAPI as a viability dye was detected with the 405nm laser and a 448/59 band pass filter. Cells were collected into Eppendorf DNA LoBind tubes containing lysis buffer, and RNA isolated using a Qiagen RNeasy Mini kit. RNA quality was assessed using RNA pico chips on an Agilent 2100 Bioanalyzer. Library preparation (SMARTer Ultra Low Input RNA for Illumina Sequencing - HV kit) and sequencing (Illumina HiSeq4000) were performed by the High-Throughput Genomics Group at the Wellcome Trust Centre for Human Genetics. The samples were processed sequenced in two batches. Batch 1 contained three control and five hypoxia samples, and batch 2 contained two repeated control samples with five ID samples.

### RNA-Seq data analysis

QC of the raw sequencing reads was performed using FastQC (https://www.bioinformatics.babraham.ac.uk/projects/fastqc). Reads were aligned to Mus musculus genome (mm10) using the splice-aware algorithm STAR v2.5.3a^98^. Gencode version M12 (Ensembl 87) was used for the annotation of the mouse genome. The DNA sequence of the GFP vector was included in the mm10 reference and its transcript was taken into account during alignment. The R Package Rsubread^99^ was used to assign and quantify the reads corresponding to each genomic feature indicated by the mm10 Gencode transcripts. Reads were assigned to the target that has the largest number of overlapping bases. The minimum fraction of overlapping bases in a read that is required for read assignment was 0.25. Counts per million (CPM) values were calculated for each sample and genes were excluded from the analysis if they did not have a CPM of a least 0.5 in a least 2 libraries. Normalisation was performed using trimmed mean of values (TMM) as implemented in the edgeR R package^100^ to scale the raw library sizes^101^. Voom transformation was performed to prepare the data for linear modelling using limma^102,103^. Principal Component Analysis and Multidimensional scaling was performed for quality control to investigate sample clustering. Batch correction was performed using the ComBat method^104^ and the sva R package^105^. Differential expression analysis between control and treated samples was performed using limma^103^. Multiple testing correction was performed on the p-values using the Benjamini-Hochberg false-discovery rate (FDR) procedure with an FDR <5%^106^. Differentially expressed genes were determined using adjusted p-value <0.01 and logFC >1 or <-1 and B >1. Unsupervised Hierarchical clustering (Extended Figure 3b) was performed using the Euclidian distance measure and complete agglomeration method and the pheatmap R package (https://cran.r-project.org/web/packages/pheatmap/index.html).

### Immunohistochemistry and X-gal staining

Immunohistochemistry on paraffin sections was done as previously described^5^. Briefly, embryos were fixed overnight in 4% paraformaldehyde at 4°C, paraffin embedded and sectioned in the indicated plane. To minimise inter-slide staining variation, tissue arrays were made by putting single sections from 12-20 different embryos on a single slide, and slides were processed using a Shandon Sequenza® Immunostaining Center (Thermo Fisher Scientific). For immunohistochemistry on wholemount hearts, tissue was fixed overnight in paraformaldehyde at 4°C. Hearts were washed three times in phosphate buffered saline plus 0.1% Triton X-100 (PBST), blocked in 5% goat serum in PBST for one hour at 4°C, then incubated in Armenian hamster anti-CD31 monoclonal antibody in PBST plus 1% bovine serum albumin overnight at 4°C. Hearts were washed three times in PBST, then incubated in biotinylated goat anti-Armenian hamster secondary antibody for 60 minutes at 4°C. Hearts were washed three times in PBST, then incubated in avidin-biotin horseradish peroxidase complex (ABC Elite, Vector PK-4000) 1:50 in PBS for 30 minutes. Finally, hearts were washed three times in PBST and placed in DAB substrate (Vector peroxidase substrate kit, SK-4100). When staining was complete, the reaction was stopped by washing in MilliQ water. X-gal staining of embryos in wholemount was done as previously described^107^.

### Antibodies

| Target | Name | Catalog number | Species and type | Supplier | Dilution |
| --- | --- | --- | --- | --- | --- |
| **Primary antibodies** |  |  |  |  |  |
| MYOSIN II HEAVY CHAIN | MF 20 |  | mouse monoclonal | Developmental Studies Hybridoma Bank<sup>108</sup> | 1:100 |
| phospho-HISTONE H3 | (Ser 10)-R | sc-8656 | rabbit polyclonal | Santa Cruz Biotechnology | 1:200 |
| GATA4 |  | sc-25310 | mouse monoclonal | Santa Cruz Biotechnology | 1:100 |
| TBX5 |  | sc-515536 | mouse polyclonal | Santa Cruz Biotechnology | 1:500 |
| NKX2-5 | N-19 | sc-8697 | goat polyclonal | Santa Cruz Biotechnology | 1:250 |
| CD31 (heart) |  | ab119341 | Armenian hamster monoclonal | Abcam | 1:500 |
| CD31 (back skin) |  | 553370 | rat monoclonal | BD Pharmingen | 1:500 |
| PROX1 |  | AF2727 | goat polyclonal | R&D systems | 1:500 |
| NRP2 | D39A5 | 3366 | rabbit monoclonal | Cell Signaling Technology | 1:500 |
| **Secondary antibodies** |  |  |  |  |  |
| Donkey anti-mouse Cy<sup>TM</sup>3 |  | 718-064-151 |  | Jackson ImmunoResearch | 1:500 |
| Donkey anti-mouse AlexaFluor® 488 |  | 715-545-151 |  | Jackson ImmunoResearch | 1:500 |
| Donkey anti-rabbit Cy<sup>TM</sup>3 |  | 711-165-152 |  | Jackson ImmunoResearch | 1:500 |
| Donkey anti-rabbit AlexaFluor® 488 |  | 711-545-152 |  | Jackson ImmunoResearch | 1:500 |
| Donkey anti-goat AlexaFluor® 488 |  | A-11055 |  | Life Technologies | 1:500 |
| Donkey anti-rat Dylight<sup>TM</sup> 405 |  | 712-475-153 |  | Jackson ImmunoResearch | 1:500 |
| Goat anti-Armenian hamster Biotinylated |  | ab5744 |  | Abcam | 1:250 |
| **Nuclear stain** |  |  |  |  |  |
| TO-PRO®-3 Iodide |  | T3605 |  | Life Technologies | 1:10,000 |

### Statistical analyses

All statistical analyses were performed with Prism 8.4.2 (GraphPad Software). Data were first tested for normal distribution by Shapiro-Wilk test and equal variance by F test. For data with two groups, normally distributed samples were tested for statistical significance using two-tailed Student’s t test (if variances equal) or two-tailed Welch’s corrected t test (if unequal variances). Non-normally distributed samples were tested using a two-tailed Mann-Whitney U test. For data with more than two groups, normally distributed data were tested using ANOVA followed by Tukey’s post-hoc test to compare the means of each group with every other group. Non-normally distributed data were tested using Kruskal–Wallis one-way ANOVA with Dunnett’s post-hoc test. The statistical significance of binomial prevalence data was tested using one-tailed Fisher’s exact test. Data are presented as mean ± standard deviation, except for Extended Figure 1 panel i, which shows median ± 95% confidence interval.

## Equipment and settings

Figure 1. Panels a,b: images were taken at 0.75x magnification on a Leica M80 dissecting microscope with a Plan 1.0x M-series objective, 0.8x video adapter, a Leica DMC4500 camera and Leica LAS software. Images were captured at 1920×2560 pixel resolution in 8 bit RGB. Adobe Photoshop was used to adjust the highlight levels to 120 equally for both images, and the Auto Color function was used on the image in panel b. Panels d-g: images were taken at 6x magnification on a Leica M80 dissecting microscope with a Plan 1.0x M-series objective, 0.8x video adapter, a Leica DMC4500 camera and Leica LAS software. Images were captured at 1920×2560 pixel resolution in 8 bit RGB. Highlight levels were adjusted to 140, and lowlight levels to 50, equally on all images in Adobe Photoshop.

Figure 3. Panels a,b,a’,b’,f,g,f’,g’: images were taken on an Olympus FV1000 confocal microscope using an Olympus UPLSAPO NA 0.75 20x objective and Fluoview FV31S-SW. Images were captured at 1024×1024 pixel resolution in 16 bit grayscale. Panels j,k: images were taken at 14x magnification on a Leica DM6000B microscope with a 20x HCX PL FLUOTAR NA 0.5 objective, a 0.7x phototube, a Leica DFC550 camera and Leica LAS software. Images were captured at 2720×2048 pixel resolution in 16 bit RGB, and cropped to 2048×2048 pixels.

Figure 4. Panels a,b: 5 image Z-stacks of tiled images (7×3 for panel 1 and 6×3 for panel b) were taken on an Olympus FV3000 confocal microscope using an Olympus UPLSAPO NA 0.4 10x objective, stitched automatically using Fluoview FV31S-SW software, and converted to a maximum intensity projection. Individual images were captured at 1024×1024 pixel resolution in 16 bit grayscale. The total size of the tiled images was 6862 × 2970 (panel a) and 5889×2970 (panel b) pixels. The image in panel a was cropped to 5889×2970 pixels. For the CD31 layers, highlight levels were adjusted to 150, and lowlight levels to 20, equally for both images in Adobe Photoshop. Panels a’-b’’’: images from panels a,b were cropped to 1000×1000 pixels. Panels e-h: images were taken at 1.75x (panels e,g) or 14x (panels f,h) magnification on a Leica DM6000B microscope with a 2.5x PL FLUOTAR NA 0.07 or a 20x HCX PL FLUOTAR NA 0.5 objective, a 0.7x phototube, a Leica DMC550 camera and Leica LAS software. Images were captured at 2720×2048 pixel resolution in 16 bit RGB. For panels e,g highlight levels were adjusted to 225, and lowlight levels to 90, equally on both images; for panels f,h highlight levels were adjusted to 225, and lowlight levels to 110, equally on all images in Adobe Photoshop. Panels i-l: images were taken at 2.5x magnification on a Leica M80 dissecting microscope with a Plan 1.0x M-series objective, 0.8x video adapter, a Leica DMC4500 camera and Leica LAS software. Images were captured at 1920×2560 pixel resolution in 8 bit RGB, and cropped to 1920×1920 pixels. Panels m,o: 10 image Z-stacks were taken on an Olympus FV3000 confocal microscope using an Olympus UPLSAPO NA 0.16 4x objective and Fluoview FV31S-SW. Images were captured at 1024×1024 pixel resolution in 16 bit grayscale. Highlight brightness was adjusted using FIJI software equally in both images. Panels n,p: 10 image Z-stacks were taken on an Olympus FV3000 confocal microscope using an Olympus UPLSAPO NA 0.95 40x objective and Fluoview FV31S-SW. Images were captured at 1024×1024 pixel resolution in 16 bit grayscale. Highlight brightness was adjusted using FIJI software equally in both images.

Figure 5. Panels c-f: images were taken at 0.75x magnification on a Leica M80 dissecting microscope with a Plan 1.0x M-series objective, 0.8x video adapter, a Leica DMC4500 camera and Leica LAS software. Images were captured at 1920×2560 pixel resolution in 8 bit RGB.

Extended Figure 1. Panels c,d: images were taken at 20x magnification using a Nikon Coolscope microscope slide scanner at 1280×960 pixel resolution in 8 bit RGB, and cropped to 960×960 pixels.

Extended Figure 3. Panel a: image was captured at 8x magnification using a Zeiss Discovery V8 microscope with PentaFluar S, a Achromat S 1.0x objective, 60N-C 1” 1.0x adapter, Axiocam 506m camera and Zeiss Zen software. The image was captured at 2752×2208 pixel resolution in 16 bit grayscale. The image was cropped to 1756×1498 pixels, highlight levels adjusted to 70 and lowlight levels adjusted to 10 using Adobe Photoshop.

Extended Figure 4. Panels a,b,e,f: images were taken on an Olympus FV1000 confocal microscope using an Olympus UPLSAPO NA 0.75 20x objective and Fluoview FV31S-SW. Images were captured at 1024×1024 pixel resolution in 16 bit grayscale. For panels e and f, highlight levels were adjusted to 140, and lowlight levels to 20, equally on both images in Adobe Photoshop. Panels h,I: images were taken at 1.25x magnification on a Leica M80 dissecting microscope with a Plan 1.0x M-series objective, 0.8x video adapter, a Leica DMC4500 camera and Leica LAS software. Images were captured at 1920×2560 pixel resolution in 8 bit RGB, cropped to 800×1500 pixels and brightness adjusted to +60 equally in both images in Adobe Photoshop.

## Data availability

The RNA-Seq data supporting the findings of this study have been deposited in the Sequence Read Archive (SRA) with BioProject ID PRJNA596545. Other data that support the findings of this study are available from the corresponding author upon reasonable request.

## Acknowledgements

This work was supported by a University of Oxford, Medical Sciences Division, Medical Research Fund grant MRF/TT2016/2207 (DBS); the Federated Foundation (DBS); an Oxford British Heart Foundation Centre of Research Excellence Senior Transition Research Fellowship RE/13/1/30181 (DBS); a BHF Senior Basic Science Research Fellowship FS/17/55/33100 (DBS); Oxford BHF CRE core infrastructure grants (DBS and JIKS, RE/18/3/34214); the John Fell Oxford University Press Research Fund (DBS); Nuffield Benefaction for Medicine and the Wellcome Trust Institutional Strategic Support Fund (DBS and NV); Novo Nordisk postdoctoral fellowships (NN, JJM); a BHF Intermediate Basic Science Research Fellowship FS/12/63/29895 (SLL); Vifor Pharma (SLL); the Francis Crick Institute, which receives its core funding from Cancer Research UK (FC001157, FC001117), the UK Medical Research Council (FC001157, FC001117) and the Wellcome Trust (FC001157, FC001117) (TJM); the Francis Crick Institute which receives its core funding from Cancer Research UK (FC001194), the UK Medical Research Council (FC001194) and the Wellcome Trust (FC001194) (VLJT); a Wellcome Trust grant 098327 (EMCF); a Wellcome Trust grant 098328 (EL-E, RA, VLJT); a Medical Research Council grant U117562103 (TJM); a National Heart Foundation of Australia Future Leader Fellowship 101204 (EG); and a NSW Health Early-Mid Career Fellowship (EG). We thank BMS staff for animal husbandry; University of Oxford Department of Earth Sciences for ICP-MS; the High-Throughput Genomics Group at the Wellcome Trust Centre for Human Genetics (funded by Wellcome Trust grant 090532/Z/09/Z) for generation of the RNA-Seq data; Michal Maj at the University of Oxford Sir William Dunn School of Pathology flow cytometry facility for sorting cells; Nadia Halidi and the Micron Advanced Bioimaging Unit (supported by Wellcome Strategic Awards 091911/B/10/Z and 107457/Z/15/Z); Ken Chien and Antonio Baldini for mouse strains; Elizabeth Robertson for use of the Leica DM6000B microscope; Simon Bamforth for help with aortic arch artery 3D modelling; and Robert Kelly and Paul Riley for critical comments on the text. The monoclonal antibody MF20^108^ developed by DA Fischman (Weill Cornell Medical College) was obtained from the Developmental Studies Hybridoma Bank, created by the NICHD of the NIH and maintained at The University of Iowa, Department of Biology, Iowa City, IA 52242.

## Author contributions (CRediT taxonomy)

Conceptualization: DBS; Methodology: JIK, NV, JJM, TJM, EG, DBS; Software JM, MT, JJM, EG; Formal analysis JIK, NV, DS, JM, MT, SEH, EG, DBS; Investigation JIK, NV, DS, JM, MT, SEH, JJM, AJ, EMS, MW, EH, FP, EL-E, RA, EG, DBS; Resources EMCF, VLJT; Writing – original draft preparation DBS; Writing – review and editing JIK, NV, DS, TJM, DBS; Visualization JIK, NV, EG, DBS; Supervision NV, VLJT, TJM, SLL, EG, DBS; Project administration DBS; Funding acquisition JIK, NV, JMM, EMCF, VLJT, TJM, SLL, EG, DBS.

## References

1 Liu, Y. et al. Global birth prevalence of congenital heart defects 1970-2017: updated systematic review and meta-analysis of 260 studies. Int J Epidemiol, doi:10.1093/ije/dyz009 (2019).

2 Alankarage, D. et al. Identification of clinically actionable variants from genome sequencing of families with congenital heart disease. Genet Med 21, 1111–1120, doi:10.1038/s41436-018-0296-x (2018).

3 Kalisch-Smith, J. I., Ved, N. & Sparrow, D. B. in Heart Development and Disease (eds B. G. Bruneau & P. R. Riley) (Cold Spring Harbor Laboratory Press, in press).

4 Worl Health Organization. The global prevalence of anaemia in 2011. (2015).

5 Shi, H. et al. Gestational stress induces the unfolded protein response, resulting in heart defects. Development 143, 2561–2572, doi:10.1242/dev.136820 (2016).

6 Yuan, X. et al. Disruption of spatiotemporal hypoxic signaling causes congenital heart disease in mice. J Clin Invest 127, 2235–2248, doi:10.1172/JCI88725 (2017).

7 Clark, R. L. et al. Diflunisal-induced maternal anemia as a cause of teratogenicity in rabbits. Teratology 30, 319–332, doi:10.1002/tera.1420300304 (1984).

8 Shepard, T. H., Mackler, B. & Finch, C. A. Reproductive studies in the iron-deficient rat. Teratology 22, 329–334, doi:10.1002/tera.1420220310 (1980).

9 Yang, J. et al. Maternal iron intake during pregnancy and birth outcomes: a cross-sectional study in Northwest China. Br J Nutr 117, 862–871, doi:10.1017/S0007114517000691 (2017).

10 Yang, J. et al. Iron intake and iron status during pregnancy and risk of congenital heart defects: A case-control study. Int J Cardiol 301, 74–79, doi:10.1016/j.ijcard.2019.11.115 (2020).

11 Chung, Y. J. et al. Iron-deficiency anemia reduces cardiac contraction by downregulating RyR2 channels and suppressing SERCA pump activity. JCI Insight 4, doi:10.1172/jci.insight.125618 (2019).

12 Francou, A., Saint-Michel, E., Mesbah, K. & Kelly, R. G. TBX1 regulates epithelial polarity and dynamic basal filopodia in the second heart field. Development 141, 4320–4331, doi:10.1242/dev.115022 (2014).

13 Rochais, F. et al. Hes1 is expressed in the second heart field and is required for outflow tract development. PLoS One 4, e6267, doi:10.1371/journal.pone.0006267 (2009).

14 Roux, M., Laforest, B., Capecchi, M., Bertrand, N. & Zaffran, S. Hoxb1 regulates proliferation and differentiation of second heart field progenitors in pharyngeal mesoderm and genetically interacts with Hoxa1 during cardiac outflow tract development. Dev Biol 406, 247–258, doi:10.1016/j.ydbio.2015.08.015 (2015).

15 MacGrogan, D. et al. How to make a heart valve: from embryonic development to bioengineering of living valve substitutes. Cold Spring Harb Perspect Med 4, a013912, doi:10.1101/cshperspect.a013912 (2014).

16 Anderson, R. H., Mori, S., Spicer, D. E., Brown, N. A. & Mohun, T. J. Development and Morphology of the Ventricular Outflow Tracts. World J Pediatr Congenit Heart Surg 7, 561–577, doi:10.1177/2150135116651114 (2016).

17 Anderson, R. H., Spicer, D. E., Mohun, T. J., Hikspoors, J. & Lamers, W. H. Remodeling of the Embryonic Interventricular Communication in Regard to the Description and Classification of Ventricular Septal Defects. Anat Rec (Hoboken) 302, 19–31, doi:10.1002/ar.24020 (2019).

18 Anderson, R. H., Brown, N. A. & Mohun, T. J. Insights regarding the normal and abnormal formation of the atrial and ventricular septal structures. Clin Anat 29, 290–304, doi:10.1002/ca.22627 (2016).

19 Gupta, S. K., Bamforth, S. D. & Anderson, R. H. How frequent is the fifth arch artery? Cardiol Young 25, 628–646, doi:10.1017/S1047951114002182 (2015).

20 Lindsay, E. A. & Baldini, A. Recovery from arterial growth delay reduces penetrance of cardiovascular defects in mice deleted for the DiGeorge syndrome region. Hum Mol Genet 10, 997–1002, doi:10.1093/hmg/10.9.997 (2001).

21 Phillips, H. M. et al. Pax9 is required for cardiovascular development and interacts with Tbx1 in the pharyngeal endoderm to control 4th pharyngeal arch artery morphogenesis. Development 146, doi:UNSP dev177618, doi:10.1242/dev.177618 (2019).

22 Priya, S., Thomas, R., Nagpal, P., Sharma, A. & Steigner, M. Congenital anomalies of the aortic arch. Cardiovasc Diagn Ther 8, S26–S44, doi:10.21037/cdt.2017.10.15 (2018).

23 Eley, L. et al. A novel source of arterial valve cells linked to bicuspid aortic valve without raphe in mice. Elife 7, doi:10.7554/eLife.34110 (2018).

24 Mifflin, J. J., Dupuis, L. E., Alcala, N. E., Russell, L. G. & Kern, C. B. Intercalated cushion cells within the cardiac outflow tract are derived from the myocardial troponin T type 2 (Tnnt2) Cre lineage. Dev Dyn 247, 1005–1017, doi:10.1002/dvdy.24641 (2018).

25 Peterson, J. C. et al. Bicuspid aortic valve formation: Nos3 mutation leads to abnormal lineage patterning of neural crest cells and the second heart field. Dis Model Mech 11, doi:10.1242/dmm.034637 (2018).

26 Liao, J. et al. Identification of downstream genetic pathways of Tbx1 in the second heart field. Dev Biol 316, 524–537, doi:10.1016/j.ydbio.2008.01.037 (2008).

27 Park, E. J. et al. An FGF autocrine loop initiated in second heart field mesoderm regulates morphogenesis at the arterial pole of the heart. Development 135, 3599–3610, doi:10.1242/dev.025437 (2008).

28 Zhang, J. et al. Frs2alpha-deficiency in cardiac progenitors disrupts a subset of FGF signals required for outflow tract morphogenesis. Development 135, 3611–3622, doi:10.1242/dev.025361 (2008).

29 Ramsbottom, S. A. et al. Vangl2-regulated polarisation of second heart field-derived cells is required for outflow tract lengthening during cardiac development. PLoS Genet 10, e1004871, doi:10.1371/journal.pgen.1004871 (2014).

30 Qyang, Y. et al. The renewal and differentiation of Isl1+ cardiovascular progenitors are controlled by a Wnt/beta-catenin pathway. Cell Stem Cell 1, 165–179, doi:10.1016/j.stem.2007.05.018 (2007).

31 Mootha, V. K. et al. PGC-1alpha-responsive genes involved in oxidative phosphorylation are coordinately downregulated in human diabetes. Nat Genet 34, 267–273, doi:10.1038/ng1180 (2003).

32 Subramanian, A. et al. Gene set enrichment analysis: a knowledge-based approach for interpreting genome-wide expression profiles. Proc Natl Acad Sci U S A 102, 15545–15550, doi:10.1073/pnas.0506580102 (2005).

33 de Soysa, T. Y. et al. Single-cell analysis of cardiogenesis reveals basis for organ-level developmental defects. Nature 572, 120–124, doi:10.1038/s41586-019-1414-x (2019).

34 Chen, L., Fulcoli, F. G., Tang, S. & Baldini, A. Tbx1 regulates proliferation and differentiation of multipotent heart progenitors. Circ Res 105, 842–851, doi:10.1161/CIRCRESAHA.109.200295 (2009).

35 Ilagan, R. et al. Fgf8 is required for anterior heart field development. Development 133, 2435–2445, doi:10.1242/dev.02408 (2006).

36 Xu, H. et al. Tbx1 has a dual role in the morphogenesis of the cardiac outflow tract. Development 131, 3217–3227, doi:10.1242/dev.01174 (2004).

37 Nemer, G. & Nemer, M. in Heart Development and Regeneration Vol. 1 (eds N. Rosenthal & R.P. Harvey) Ch. 9.2, 599–616 (Academic Press, 2010).

38 Billings, S. E. et al. The retinaldehyde reductase DHRS3 is essential for preventing the formation of excess retinoic acid during embryonic development. FASEB J 27, 4877–4889, doi:10.1096/fj.13-227967 (2013).

39 Bowles, J. et al. Control of retinoid levels by CYP26B1 is important for lymphatic vascular development in the mouse embryo. Developmental Biology 386, 25–33, doi:10.1016/j.ydbio.2013.12.008 (2014).

40 Roberts, C., Ivins, S., Cook, A. C., Baldini, A. & Scambler, P. J. Cyp26 genes a1, b1 and c1 are down-regulated in Tbx1 null mice and inhibition of Cyp26 enzyme function produces a phenocopy of DiGeorge Syndrome in the chick. Hum Mol Genet 15, 3394–3410, doi:10.1093/hmg/ddl416 (2006).

41 Cohlan, S. Q. Excessive intake of vitamin A as a cause of congenital anomalies in the rat. Science 117, 535–536, doi:10.1126/science.117.3046.535 (1953).

42 Lammer, E. J. et al. Retinoic Acid Embryopathy. New Engl J Med 313, 837–841, doi:Doi 10.1056/Nejm198510033131401 (1985).

43 Arceci, R. J., King, A. A., Simon, M. C., Orkin, S. H. & Wilson, D. B. Mouse GATA-4: a retinoic acid-inducible GATA-binding transcription factor expressed in endodermally derived tissues and heart. Mol Cell Biol 13, 2235–2246, doi:10.1128/mcb.13.4.2235 (1993).

44 Ghatpande, S., Ghatpande, A., Zile, M. & Evans, T. Anterior endoderm is sufficient to rescue foregut apoptosis and heart tube morphogenesis in an embryo lacking retinoic acid. Dev Biol 219, 59–70, doi:10.1006/dbio.1999.9601 (2000).

45 Rossant, J., Zirngibl, R., Cado, D., Shago, M. & Giguere, V. Expression of a Retinoic Acid Response Element-Hsplacz Transgene Defines Specific Domains of Transcriptional Activity during Mouse Embryogenesis. Gene Dev 5, 1333–1344, doi:DOI 10.1101/gad.5.8.1333 (1991).

46 Ghyselinck, N. B. & Duester, G. Retinoic acid signaling pathways. Development 146, doi:10.1242/dev.167502 (2019).

47 Wilson, R. et al. Highly variable penetrance of abnormal phenotypes in embryonic lethal knockout mice. Wellcome Open Res 1, 1, doi:10.12688/wellcomeopenres.9899.2 (2016).

48 Bellini, C. et al. Etiology of non-immune hydrops fetalis: An update. Am J Med Genet A 167A, 1082–1088, doi:10.1002/ajmg.a.36988 (2015).

49 Betterman, K. L. & Harvey, N. L. Histological and Morphological Characterization of Developing Dermal Lymphatic Vessels. Methods Mol Biol 1846, 19–35, doi:10.1007/978-1-4939-8712-2_2 (2018).

50 Qu, X., Tompkins, K., Batts, L. E., Puri, M. & Baldwin, H. S. Abnormal embryonic lymphatic vessel development in Tie1 hypomorphic mice. Development 137, 1285–1295, doi:10.1242/dev.043380 (2010).

51 Sharma, B., Chang, A. & Red-Horse, K. Coronary Artery Development: Progenitor Cells and Differentiation Pathways. Annu Rev Physiol 79, 1–19, doi:10.1146/annurev-physiol-022516-033953 (2017).

52 Chen, H. I. et al. The sinus venosus contributes to coronary vasculature through VEGFC-stimulated angiogenesis. Development 141, 4500–4512, doi:10.1242/dev.113639 (2014).

53 Tian, X. Y., Pu, W. T. & Zhou, B. Cellular Origin and Developmental Program of Coronary Angiogenesis. Circulation Research 116, 515–530, doi:10.1161/Circresaha.116.305097 (2015).

54 Wessels, A. et al. Epicardially derived fibroblasts preferentially contribute to the parietal leaflets of the atrioventricular valves in the murine heart. Dev Biol 366, 111–124, doi:10.1016/j.ydbio.2012.04.020 (2012).

55 Smart, N. et al. Thymosin beta4 induces adult epicardial progenitor mobilization and neovascularization. Nature 445, 177–182, doi:10.1038/nature05383 (2007).

56 Wang, S. et al. Alterations in retinoic acid signaling affect the development of the mouse coronary vasculature. Dev Dyn 247, 976–991, doi:10.1002/dvdy.24639 (2018).

57 Fernandez, B. et al. The coronary arteries of the C57BL/6 mouse strains: implications for comparison with mutant models. J Anat 212, 12–18, doi:10.1111/j.1469-7580.2007.00838.x (2008).

58 Wheby, M. S. & Crosby, W. H. The Gastrointestinal Tract and Iron Absorption. Blood 22, 416–428 (1963).

59 Santos, M. et al. In vivo mucosal uptake, mucosal transfer and retention of iron in mice. Lab Anim 31, 264–270, doi:10.1258/002367797780596329 (1997).

60 Sangkhae, V. et al. Effects of maternal iron status on placental and fetal iron homeostasis. J Clin Invest, doi:10.1172/JCI127341 (2019).

61 Sakabe, M., Kokubo, H., Nakajima, Y. & Saga, Y. Ectopic retinoic acid signaling affects outflow tract cushion development through suppression of the myocardial Tbx2-Tgfbeta2 pathway. Development 139, 385–395, doi:10.1242/dev.067058 (2012).

62 Yasui, H., Morishima, M., Nakazawa, M., Ando, M. & Aikawa, E. Developmental spectrum of cardiac outflow tract anomalies encompassing transposition of the great arteries and dextroposition of the aorta: pathogenic effect of extrinsic retinoic acid in the mouse embryo. Anat Rec 254, 253–260, doi:10.1002/(SICI)1097-0185(19990201)254:2<253::AID-AR11>3.0.CO;2-4 (1999).

63 Yasui, H., Nakazawa, M., Morishima, M., Miyagawa-Tomita, S. & Momma, K. Morphological observations on the pathogenetic process of transposition of the great arteries induced by retinoic acid in mice. Circulation 91, 2478–2486, doi:10.1161/01.cir.91.9.2478 (1995).

64 Niderla-BieliNska, J. et al. Proepicardium: Current Understanding of its Structure, Induction, and Fate. Anat Rec (Hoboken) 302, 893–903, doi:10.1002/ar.24028 (2019).

65 Bautch, V. L. & Caron, K. M. Blood and lymphatic vessel formation. Cold Spring Harb Perspect Biol 7, a008268, doi:10.1101/cshperspect.a008268 (2015).

66 Mason, K. E. Foetal death, prolonged gestation, and difficult parturition in the rat as a result of Vitamin A-deficiency. Am J Anat 57, 303–349 (1935).

67 Prendiville, T., Jay, P. Y. & Pu, W. T. Insights into the genetic structure of congenital heart disease from human and murine studies on monogenic disorders. Cold Spring Harb Perspect Med 4, doi:10.1101/cshperspect.a013946 (2014).

68 Chapman, G. et al. Functional genomics and gene-environment interaction highlight the complexity of Congenital Heart Disease caused by Notch pathway variants. Hum Mol Genet, doi:10.1093/hmg/ddz270 (2019).

69 Moreau, J. L. M. et al. Gene-environment interaction impacts on heart development and embryo survival. Development 146, doi:10.1242/dev.172957 (2019).

70 Morrow, B. E., McDonald-McGinn, D. M., Emanuel, B. S., Vermeesch, J. R. & Scambler, P. J. Molecular genetics of 22q11.2 deletion syndrome. Am J Med Genet A 176, 2070–2081, doi:10.1002/ajmg.a.40504 (2018).

71 Hartman, R. J. et al. The contribution of chromosomal abnormalities to congenital heart defects: a population-based study. Pediatr Cardiol 32, 1147–1157, doi:10.1007/s00246-011-0034-5 (2011).

72 Lindsay, E. A. et al. Tbx1 haploinsufficieny in the DiGeorge syndrome region causes aortic arch defects in mice. Nature 410, 97–101, doi:10.1038/35065105 (2001).

73 Lana-Elola, E. et al. Genetic dissection of Down syndrome-associated congenital heart defects using a new mouse mapping panel. Elife 5, doi:10.7554/eLife.11614 (2016).

74 Chou, H. H. et al. Association of maternal chronic disease with risk of congenital heart disease in offspring. CMAJ 188, E438–E446, doi:10.1503/cmaj.160061 (2016).

75 Liu, S. et al. Association between maternal chronic conditions and congenital heart defects: a population-based cohort study. Circulation 128, 583–589, doi:10.1161/CIRCULATIONAHA.112.001054 (2013).

76 Robinson, R. et al. Risk factors for congenital heart defects in two populations residing in the same geographic area: a long-term population-based study, Southern Israel. Cardiol Young, 1–5, doi:10.1017/S1047951119001409 (2019).

77 Kolodzinska, A., Heleniak, A. & Ratajska, A. Retinoic acid-induced ventricular non-compacted cardiomyopathy in mice. Kardiol Pol 71, 447–452, doi:10.5603/KP.2013.0090 (2013).

78 Andreini, C., Putignano, V., Rosato, A. & Banci, L. The human iron-proteome. Metallomics 10, 1223–1231, doi:10.1039/c8mt00146d (2018).

79 Cuny, H. et al. NAD deficiency due to environmental factors or gene-environment interactions causes congenital malformations and miscarriage in mice. Proc Natl Acad Sci U S A 117, 3738–3747, doi:10.1073/pnas.1916588117 (2020).

80 Shi, H. et al. NAD Deficiency, Congenital Malformations, and Niacin Supplementation. N Engl J Med 377, 544–552, doi:10.1056/NEJMoa1616361 (2017).

81 Karczewski, K. J. et al. The mutational constraint spectrum quantified from variation in 141,456 humans. Nature 581, 434–443, doi:10.1038/s41586-020-2308-7 (2020).

82 Maclean, G., Dolle, P. & Petkovich, M. Genetic disruption of CYP26B1 severely affects development of neural crest derived head structures, but does not compromise hindbrain patterning. Dev Dyn 238, 732–745, doi:10.1002/dvdy.21878 (2009).

83 Okano, J. et al. The regulation of endogenous retinoic acid level through CYP26B1 is required for elevation of palatal shelves. Dev Dyn 241, 1744–1756, doi:10.1002/dvdy.23862 (2012).

84 Kalter, H. & Warkany, J. Experimental production of congenital malformations in strains of inbred mice by maternal treatment with hypervitaminosis A. Am J Pathol 38, 1–21 (1961).

85 Kallen, B., Mastroiacovo, P. & Robert, E. Major congenital malformations in Down syndrome. Am J Med Genet 65, 160–166, doi:10.1002/(SICI)1096-8628(19961016)65:2<160::AID-AJMG16>3.0.CO;2-O (1996).

86 Okubo, T. et al. Ripply3, a Tbx1 repressor, is required for development of the pharyngeal apparatus and its derivatives in mice. Development 138, 339–348, doi:10.1242/dev.054056 (2011).

87 Janesick, A., Shiotsugu, J., Taketani, M. & Blumberg, B. RIPPLY3 is a retinoic acid-inducible repressor required for setting the borders of the pre-placodal ectoderm. Development 139, 1213–1224, doi:10.1242/dev.071456 (2012).

88 Jerome, L. A. & Papaioannou, V. E. DiGeorge syndrome phenotype in mice mutant for the T-box gene, Tbx1. Nat Genet 27, 286–291, doi:10.1038/85845 (2001).

89 Pierson, J. C., Ferris, L. K. & Schwarz, E. B. We Pledge to Change iPLEDGE. JAMA Dermatol 151, 701–702, doi:10.1001/jamadermatol.2015.0736 (2015).

90 World Health Organization. Global prevalence of vitamin A deficiency in populations at risk 1995–2005. WHO Global Database on Vitamin A Deficiency. (World Health Organization, 2009).

91 Pavord, S. et al. UK guidelines on the management of iron deficiency in pregnancy. Br J Haematol 156, 588–600, doi:10.1111/j.1365-2141.2011.09012.x (2012).

92 World Health Organization. Guideline: Daily iron and folic acid supplementation in pregnant women. Geneva (2012).

93 Lakhal-Littleton, S. et al. Cardiac ferroportin regulates cellular iron homeostasis and is important for cardiac function. Proc Natl Acad Sci U S A 112, 3164–3169, doi:10.1073/pnas.1422373112 (2015).

94 Cleary, J. O. et al. Cardiac phenotyping in ex vivo murine embryos using microMRI. NMR Biomed 22, 857–866, doi:10.1002/nbm.1400 (2009).

95 Zamyadi, M. et al. Mouse embryonic phenotyping by morphometric analysis of MR images. Physiol Genomics 42A, 89–95, doi:10.1152/physiolgenomics.00091.2010 (2010).

96 Margosian, P., DeMeester, G. & Liu, H. in eMagRes (eds R.K Harris & R.L. Wasylishen). doi: 0.1002/9780470034590.emrstm0376 (2007).

97 Weninger, W. J. & Mohun, T. J. Three-dimensional analysis of molecular signals with episcopic imaging techniques. Methods Mol Biol 411, 35–46, doi:10.1007/978-1-59745-549-7_4 (2007).

98 Dobin, A. et al. STAR: ultrafast universal RNA-seq aligner. Bioinformatics 29, 15–21, doi:10.1093/bioinformatics/bts635 (2013).

99 Liao, Y., Smyth, G. K. & Shi, W. The R package Rsubread is easier, faster, cheaper and better for alignment and quantification of RNA sequencing reads. Nucleic Acids Res 47, doi:ARTN e47, doi:10.1093/nar/gkz114 (2019).

100 McCarthy, D. J., Chen, Y. S. & Smyth, G. K. Differential expression analysis of multifactor RNA-Seq experiments with respect to biological variation. Nucleic Acids Res 40, 4288–4297, doi:10.1093/nar/gks042 (2012).

101 Robinson, M. D. & Oshlack, A. A scaling normalization method for differential expression analysis of RNA-seq data. Genome Biol 11, doi:ARTN R25, doi:10.1186/gb-2010-11-3-r25 (2010).

102 Law, C. W., Chen, Y. S., Shi, W. & Smyth, G. K. voom: precision weights unlock linear model analysis tools for RNA-seq read counts. Genome Biol 15, doi:ARTN R29, doi:10.1186/gb-2014-15-2-r29 (2014).

103 Ritchie, M. E. et al. limma powers differential expression analyses for RNA-sequencing and microarray studies. Nucleic Acids Res 43, doi:ARTN e47, doi:10.1093/nar/gkv007 (2015).

104 Johnson, W. E., Li, C. & Rabinovic, A. Adjusting batch effects in microarray expression data using empirical Bayes methods. Biostatistics 8, 118–127, doi:10.1093/biostatistics/kxj037 (2007).

105 Leek, J. T. et al. sva: Surrogate Variable Analysis. R package version 3.32.1 (2019).

106 Benjamini, Y. & Hochberg, Y. Controlling the False Discovery Rate - a Practical and Powerful Approach to Multiple Testing. J R Stat Soc B 57, 289–300 (1995).

107 Hogan, B., Beddington, R., Costantini, F. & Lacy, E. Manipulating the Mouse Embryo. A Laboratory Manual. Second edn, (Cold Spring Harbor Laboratory Press, 1994).

108 Bader, D., Masaki, T. & Fischman, D. A. Immunochemical analysis of myosin heavy chain during avian myogenesis in vivo and in vitro. J Cell Biol 95, 763–770, doi:10.1083/jcb.95.3.763 (1982).

